# The heterogeneity of the DNA damage response landscape determines patient outcomes in ovarian cancer

**DOI:** 10.1101/2021.08.03.454868

**Authors:** TDJ Walker, ZF Faraahi, MJ Price, A Hawarden, CA Waddell, B Russell, DM Jones, A McCormick, N Gavrielides, S Tyagi, LC Woodhouse, B Whalley, C Roberts, EJ Crosbie, RJ Edmondson

## Abstract

Defective DNA damage response (DDR) pathways allow cancer cells to accrue genomic aberrations and evade normal cellular growth checkpoints. Defective DDR also determines response to chemotherapy. However, the interaction and overlap between the two double strand repair pathways and the three single strand repair pathways is complex, and has remained poorly understood.

Here we show that, in ovarian cancer, a disease hallmarked by chromosomal instability, explant cultures show a range of DDR abrogation patterns. Defective homologous recombination (HR) and non-homologous end joining (NHEJ) are near mutually exclusive with HR deficient (HRD) cells showing increased abrogation of the single strand repair pathways compared to NHEJ defective cells.

When combined with global markers of DNA damage, including mitochondrial membrane functionality and reactive oxygen species burden, the pattern of DDR abrogation allows the construction of DDR signatures which are predictive of both *ex vivo* cytotoxicity, and more importantly, patient outcome.

**Significance:** Holistic assessment of the DDR is possible, shows improved ability to predict response to chemotherapy over single pathway assessment, and is applicable to a variety of ovarian cancer types. Such an assessment has clinical utility in settings of therapeutic dilemma such as retreatment for relapse.

## 1. Intro

An impaired DNA damage response (DDR) is fundamental to the development of the genome instability that defines cancer [1]. The DDR is complex with overlapping pathways but can be represented as five pathways; two for double strand breaks (DSB) and three for single strand breaks (SSB).

The role of the homologous recombination (HR) pathway is perhaps the best described of the five DDR pathways. Tasked with high-fidelity homology-based repair of DNA DSB, HR repair is closely linked to cell cycle to ensure DNA is repaired prior to mitosis or S phase [2, 3]. Non-homologous end joining (NHEJ) represents the second DSB DNA repair pathway present in the cell. NHEJ can directly ligate broken DNA and can accommodate non-compatible sequences with non-complementary break overhangs (reviewed in [4]). NHEJ operates throughout the cell cycle thus, whilst error prone, NHEJ represents the predominant DSB repair DDR pathway in human cells.

Three DDR pathways each govern distinct forms of SSB DNA repair: The base excision repair (BER) pathway responds to non-helix distorting single-base lesions such as oxidized bases, deaminated bases, and alkylated bases that are caused by reactive oxygen species (ROS) or ionizing radiation assault [5]. The nucleotide excision repair (NER) repairs helix-distorting bulky lesions and crosslinks [5, 6], and the mismatch repair (MMR) pathway responds to Watson-Crick mismatch base erroneous insertion, deletion, and mis-incorporations [7, 8].

However, the reality is that there is overlap and redundancy between DDR pathways [9]. Moreover the DDR needs to be seen in the context of other intracellular processes including mitochondrial (Mt) dysfunction and (ROS) both of which influence chemotherapy response [10]. Chemoresistance is multifaceted and driven by mechanistic and temporal factors connected to both the tumour microenvironment and the intrinsic ability of the ovarian cancer cell to resist CT [11]. Platinum CT is accepted to act by increasing intracellular ROS beyond a critical redox-homeostasis threshold from which cancer cells cannot recover [10]. Chemoresistant cancer cells likely evolve rapid homeostatic recovery or increased tolerances to accommodate the frequently-observed ROS abundance and oxidative stress increases [12] while perturbed ROS signalling enhances cell proliferation and survival [13]. Mitochondria are key producers and modulators of ROS [14] and so highly relevant to platinum CT efficacy. It is established that cancer cell Mt mutations act as oncogenomes [15] to influence oncogenesis [16] wherein chemoresistant cell Mt dysfunction associates with aggressive ovarian cancer subtypes, apoptotic resistance [17], and metastasis independent of the microenvironment [18].

Ovarian cancer is a heterogeneous set of diseases characterised by chromosomal instability making it an ideal model for profiling the DDR landscape [19]. Although individual pathway aberrations have been described for HR [20, 21] and NHEJ [22, 23] in particular, a comprehensive assessment has not been carried out.

To this end we developed a panel of *ex vivo* assays with the intention of providing comprehensive DDR pathway measures incorporating both mitochondrial dysfunction scores and ROS capacities in order to monitor the DDR landscape in ovarian cancer explant cultures. We trialled this functional DDR assay panel on explant cultures established from ovarian tumour samples from patients undergoing primary surgery or interval debulking surgery and correlated the signatures seen with clinical outcome.

## 2. Methods

### 2.1. Ethics, recruitment, and data collection

Samples were obtained from patients undergoing primary or delayed primary surgery for advanced ovarian cancer. Informed consent was obtained from all patients for sample and data collection and the study was approved by ethics.(NHS National Research Ethics Service (NRES) Committee North West approval numbers 14/NW/1260 or 19/NW/0644 depending on date of collection). Histological diagnosis was confirmed by specialist gynaecological pathologists, and patient data were obtained from patient records including time from diagnosis to first relapse (progression free survival) and time from diagnosis to death (overall survival).

### 2.2. Explant establishment and routine culture

Solid tumour specimens and ascitic fluid samples were transported fresh to the laboratory. Acellular and calcified structures in solid specimens were excised and specimens were incubated with ≥0.1U/ml Collagenase and 0.8U/ml Dispase (Roche) for 120 minutes at 37°C 5% CO_2_ as per manufacturer’s recommendations. Explants were established from resuspended solid tumour cells while ascitic cultures were directly transferred into culture media.

Standard culture media comprised RPMI adjusted to 20% bovine serum (BS) with 2mM glutamine, 120 units/ml Penicillin G Sodium, 100µg/ml Streptomycin, and 0.5µg/ml Amphotericin B (all media and supplements were Gibco). Explants were passaged at 70% confluency and cryopreservation stocks were prepared from P0 passage times in 90% BS 10% DMSO (Thermo Fisher).

### 2.3. Explant characterisation

Cells were fixed in ice-cold methanol for 30 minutes, then exposed to a 5% Goat Serum (Sigma, G9023) block at room temperature for 1 hour. Wells were then treated with one of the following antibodies: Anti-Ca125 (Abcam, ab1107), Anti-Pax8 (Abcam, ab53490), Anti-Vimentin (Abcam, ab92547), and Anti-Pan Cytokeratin (Merck, cbl234f), all at a concentration of 1:100 in 5% Goat Serum for 1 hour at room temperature.

All wells were then washed, and Ca125, Pax8 wells, and one unstained well were exposed to Goat Anti-Mouse IgG, Alexa Fluor 546 (Invitrogen, A-11003) at a concentration of 2µg/ml. The vimentin well, and one unstained well were exposed to Goat Anti-Rabbit IgG, Alexa fluor 488 (Invitrogen, A-11008) at a concentration of 2µg/ml. Both secondary antibodies were incubated for one hour at room temperature. All staining was imaged using a Zeiss Axio Observer Z1 with Zeiss Zen 2.3 software. Antibody information is provided in supplementary methods section 1.1.

### 2.4. Rad 51 assay (HR)

HR was assessed as previously described [24]. Briefly, cells were seeded at a density of 40,000 cells/cm2 and incubated for 24 hours. Cells were exposed to 200mJ/cm^2^ of UV-C radiation, incubated for 2 hours, and fixed in ice-cold methanol for 30 minutes. Following permeabilisation cells were treated with anti-phosphorylated H2AX (Millipore, 05-636-I) and anti-Rad51 (Abcam, ab133534) antibodies at concentrations of 1µg/ml and 1:1000 respectively followed by exposure to Goat Anti-Mouse IgG Alexa Fluor 546 (Invitrogen, A-11003) and Goat Anti-Rabbit IgG Alexa Fluor 488 (Invitrogen, A-11008) secondary antibodies at 2ug/ml. Staining was imaged using a Zeiss Axio Observer Z1 with Zeiss Zen 2.3 software. ImageJ software was used to identify nuclear regions using DAPI. A two-fold increase in the average number of phosphorylated H2AX foci in irradiated versus control cells demonstrated sufficient assault to the cells. Our previously validated threshold of a two-fold increase in the average number of Rad51 foci versus control signalled a competent cell sample [24]. Antibody information is provided in supplementary methods section 1.2.

### 2.5. *In vitro* cell extract assay (NHEJ)

The cell extract assay (NHEJ) used for the optimisation cohorts was carried out as previously described [22]. Further details are provided in supplementary methods section 1.3.

### 2.6. Host-cell reactivation assay (NHEJ)

The Host-cell Reactivation system proposed by Nagel *et al* [25] was expanded to provide per-cell quantitative NHEJ pathway capacity monitoring of blunt end, 5’-3’ and 3’-5’ discontiguous de-phosphorylated mismatched overhang DNA double-strand breaks. In summary XL1-Blue supercompetent cells (200236; Agilent) were transformed with reconstituted pCMV6-AC-GFP plasmid (ps100010; Origene). EcoRI-HF (R3101S), SacI-HF (R3156S), KpnI-HF (R3142S), PmeI (R0560S), and Quick CIP (M0525) (New England Biolabs) were used with NEB CutSmart buffer and nuclease-free water to construct the three dephosphorylated DSB plasmid conformations described in this project. Explant plasmid transfections were prepared with lipofectamine LTX Plus or 3000 (15338-100, 11668-027; Thermo Fisher) using Opti-Mem (11058021; Thermo Fisher) as per manufacturer’s guidelines. Cultures were monitored at 24, 48, and 72 hours prior to analysis. Fluorescence-governed NHEJ pathway activity of each sample was assessed qualitatively by microscopy and quantitatively by flow cytometry. Fluorescence change is concordant with traditional cell-free extract NHEJ assays and provides per-explant NHEJ pathway capacities. Values were standardised to parent negative and transfection positive control conditions to provide percent-scaled capacities. Cell event monitoring enables per-cell resolution of intra-explant NHEJ repair capacity variance and thus enables detection of possible basal level NHEJ activity which would traditionally be obfuscated below aggregate-assay sensitivity thresholds. The ratio of event frequency of positive threshold cells to all sample cells was calculated to provide the proportion of cells within any sample that demonstrated repair activity. The event ratio of positive control samples represents the maximum achieved transfection efficiency per explant and was used to percentage-calibrate target condition group ratios.

An integrated NHEJ capacity scoring system was derived from parent-normalised equal-weighted contributions of cell event and fluorescent intensity standardised percent values. *Ordinal plasmid condition scores* were: Fully Defective (<5%); Basal Activity (≥5% & <15%); Low Activity (≥15%& <25%); Competent Activity (≥25% & < 35%); and High Activity (≥35%). *Plasmid condition scores* were combined to provide *Explant capacity scores* which measure overall explant NHEJ repair capacity. *Ordinal explant capacity scores* provided a 12-point scale (driven from the three input five-point scales).

Detailed information is provided in supplementary methods section 1.4 and supplementary data 1.

### 2.7. Single Cell Gel Electrophoresis Assay (BER & NER)

The alkaline version of the single cell gel electrophoresis (comet) assay popularised by Singh [26] was adopted to monitor BER and NER pathway capacities. The assays employed pathway-relevant DNA damaging mechanisms and pathway-specific inhibition agents as outlined below. Both pathways were monitored concurrently to minimise variation. Each pathway assay comprised four conditions, each of which contained identical cell number and incubation volumes. BER and NER assays each comprised four incubation conditions containing 400,000 explant cells at 250,000 cells / ml. Antibiotics & antifungal were removed from cultures 24hrs prior to assay preparation.

### BER assault conditions

*Condition one:* negative control (sterile-filtered DMSO). *Condition two:* pathway repair blockade (10µM Olaparib (A10111-100, Generon Ltd)) in DMSO. *Condition three:* genomic assault (200µM H_2_O_2_ dH_2_O). *Condition four:* Damage during repair blockage (combined exposure to agents in conditions two and three). Cells were acclimatised for 60 minutes at 37°C prior to treatment with Olaparib and H_2_O_2_ for 20 minutes on ice. Samples were washed with 4°C dPBS, resuspended in 37°C standard media, and incubated for 90 minutes at 37°C 5% CO_2_.

### NER assault conditions

*Condition one:* negative control (sterile-filtered DMSO). *Condition two:* pathway repair blockade (50µg/ml aphidicolin (Tocris; CAS No: 38966-21-1)) in DMSO. *Condition three:* genomic assault (50µM Benzo[a]pyrene (B[a]P) (Sigma)) DMSO. *Condition four:* Damage during repair blockage (combined exposure to agents in conditions two and three). Cells were acclimatised for 30 minutes prior to treatment with aphidicolin. After a further 30 minutes cells were treated with B[a]P and incubated for 150 minutes at 37°C 5% CO_2_.

### Comet slide preparation

Standard microscope glass slides were coated with 1% ultrapure agarose in ultrapure water (16500500, 10977035; Thermo Fisher) and maintained in the dark once set. Cells were pelleted for three minutes at 1000rcf 4°C, washed in sterile 4°C dPBS (D8537, Sigma), pelleted again and dPBS was removed. Pellets were resuspended in 1% low melting point agarose (V2111; Promega) in dPBS, and samples were distributed across replicate comet slides and allowed to set. 0.5% LMP agarose was distributed over samples and allowed to set.

### Electrophoresis

Slides were transferred into electrophoresis tanks and submerged in fresh 4°C electrophoresis solution (200mM NaOH; 1mM Na_2_EDTA•2H_2_0 (pH 10), in distilled water, confirmed pH >13.0) and incubated for 45 minutes at 4°C in the dark to permit DNA unwinding. Electrophoresis was conducted for 36 minutes at 1.5V/cm with a 2mm buffer height above the comet slide hyperplane.

### Comet imaging and pathway competence score generation

Slides were visualised at 10x magnification using a Zeiss Axio Observer Z1 with Zeiss Zen 2.3 software. 172-megapixel tiled images were acquired and comets were scored using Trevigen Analysis Software (4260-000-CS; Trevigen). Between 6,000 and 14,000 comets were routinely scored per each of the four explant conditions for BER or NER assays. The lowest 10 percentiles per each explant assay condition were removed to reduce uninformative values and the median percent DNA in tail per each condition was determined. DNA quantitative capacity metrics were calculated as *conditions* (4 – 1) – (3 – 1) – (2 – 1). Explant *condition three* percent DNA in tail values were summed with the median *condition two* equivalent value. These were compared with *condition four* to determine significant BER & NER pathway capacity changes (Kruskal Wallis, alpha = 0.05). Normalised survivorship class balance fold-change was calculated as the number of comets / number of *condition one* comets. Extent of percent DNA in tail variance per each condition was determined and served as a surrogate for intra-explant heterogeneity.

### 2.8. Whole Exome Sequencing (MMR)

In contrast to the other pathways which comprise multiple overlapping components, the MMR pathway can be defined by a 22 gene panel and does not require functional analysis. Purified explant DNA (QIAamp DNA Mini, 51304; Qiagen) was assessed for purity and yield (NanoDrop ND-1000). SureSelect Human All Exon V6 (Agilent) exome libraries were prepared and samples were paired-end whole exome sequenced (WES) on a DNBSEQ™ NGS Platform (BGI Genomics, Hong Kong) with a per-sample 100x depth-of-coverage target. Trimmed and cleaned reads (minimum Phred: 34.4) were aligned to human reference build Hg19 / GRCh37 using BWA [27] (99.954% average alignment) and variants were called using SAMtools [28]. Variants were annotated by dbSNP [29], SnpEff [30], and Variant Effect Predictor [31]. Outputs were recalibrated, summarised and intersect and union consensus cohorts were generated. Canonical MMR genes (*MLH1 MSH2 MSH6 PMS2*, review: [7]) and a 22-gene extended MMR panel [32] were extracted and variant mutations inspected by their type, frequency, location within the ORF, and SnpEff & VEP ontological mutation Impact annotations. Explant MMR pathway capacity scores were generated based on the frequency of mutations observed per each ontological impact level severity across the MMR gene cohort. MMR intra-gene and inter-gene mutation frequency aggregates were weighted by Impact level severities and scored ordinally as Competent; Functioning; Reduced; Perturbed; and Defective.

### 2.9. Explant platinum cytotoxicity and GR_50_ modelling

Explant cells were seeded at 2,000 cells per well in 100µl volumes of replicate 96 well culture plates. At 24 hours cells were PBS washed and triplicate wells were incubated with a 12-point scale (0µM, 1µM, 4µM, 8µM, 16µM, 32µM, 64µM, 128µM, 256µM, 512µM, 1024µM, 2048µM) of carboplatin (A10182, Generon Ltd) prepared in DMSO for 72 hours at 37°C 5% CO_2_ alongside corresponding cell-free drug with media negative controls. Cells were supplemented with 20µl of CellTiter 96® AQueous One reagent (Promega) and incubated at 37°C 5% CO_2_ in the dark. Colorimetric absorbance was measured at 450nm at time points 30, 60, 90, 120, and 180 minutes (FLUOstar Omega, BMG). Explant OD values were subtractive normalised against negative control values and intra-plate and inter-plate replicate variance was inspected. GR_50_ [33] growth-adjusted IC_50_ models were generated for each explant.

### 2.10. Intra-explant Reactive Oxygen Species (ROS) burden

The explant cell response to ROS assault was assayed using DCFDA Cellular ROS detection kits (ab113851, Abcam) with adjustments noted below. Explant cells were seeded at 25,000 cells per well and incubated with 20µM DCFDA for 45 minutes at 37°C 5% CO_2_ in the dark. Replicate wells were incubated with an 8-point scale (0µM, 4µM, 8µM, 16µM, 32µM 64µM, 128µM, 256µM) of Tert-Butyl Hydrogen Peroxide (TBHP) Plates were sealed and fluorescence was measured at Ex/Em 485/535 nm (FLUOstar Omega, BMG). Endogenous fluorescence in the presence/absence of TBHP was 0.2-0.23 fold the detection of cell with DCFDA conditions and indicative that all monitored fluorescent changes were ROS-actuated events.

### 2.11. Intra-explant Mitochondrial Membrane Function

Explant mitochondrial membrane potential was assayed using JC-10 kits (ab112134, Abcam) with adjustments noted below. Explant cells were seeded at 25,000 cells per well in 100µl volumes of clear-base black fluorescent 96 well culture plates. At 24 hours cells were washed with PBS and incubated with a 5-point scale (0mM, 0.2mM, 0.4mM, 0.8mM, 1.6mM) of H_2_O_2_ for 75 minutes at 37°C 5% CO_2_ in the dark. Wells were supplemented with 30µM JC-10 and incubated for 60 minutes at 37°C 5% CO_2_ in the dark. Assay Buffer B was added to each well and plates were sealed. Fluorescence was measured at both Ex/Em 490/525 nm and 540/590 nm (FLUOstar Omega, BMG). Intra-plate and inter-plate replicate mean and standard deviation was calculated per each drug concentration. Explant baseline mitochondrial membrane activity ratios and parent-normalised activity changes in response to the presence of drug were calculated and used to score mitochondrial status.

### 2.12. Statistical analysis, capacity scoring, classification, modelling, validation, dimension reduction

#### Software & hardware

Operating Systems: Windows 7 & 10 x64; Linux Ubuntu 14.04.6 LTS. Microsoft Excel 2016 & 2019 were used for preprocessing and collating of clinical data from NHS resources. Microsoft Access 2016 and 2019 were used for biobank-resolution (surgery-instance) databases. The R language [34] (versions 3.4.3 to 4.0.5) and RStudio [35] (1.1.456 to 1.4.1106-5) were used for explant-resolution databases and for patient-resolution databases. Over 30 packages were used in addition to *BaseR* (supplementary methods section 1.5).

#### Database handling

Bench lab data for the five DDR assays and three cytotoxicity & metabolic assays were pre-processed and contained within their own R environments. Explant-resolution summary data was migrated and mapped to both biobank-resolution idents and hashed patient NHS idents. This permitted bidirectional shuttling of data at the explant, biobank, and patient levels for classification purposes. For patient idents that mapped to multiple explant tissues, explant-resolution bench lab findings were merged to patient-resolution summaries by a framework that accommodated any discordant lab findings by weightings from significance values, quantitative capacity score, variance, class-balance, multi-tissue frequency, and extent of discordant states across all explants in question (supplementary methods section 1.5). Binomial metrics for the Rad51 (HR) assay can only by definition resolve to a summary value via odd explant numbers. Any equally weighted even number findings were ordinal-scored as heterogeneous.

#### Statistical analysis, classification, machine learning

Normality was assessed by Shapiro–Wilk or Kolmogorov–Smirnov tests and by Q-Q plots. Heteroscedasticity was determined by Bartlett’s, Levene’s, or Fligner-Killeen’s. Parametric tests included t test, ANOVA, or Welch’s ANOVA with Turkey HSD, Dunnet’s test, Bonferroni correction, or Benjamini-Hochberg’s false discovery rate post-hoc tests as appropriate. Nonparametric tests included Wilcoxon Rank Sum, Wilcoxon Signed Rank, or Kruskal Wallis tests with Dunn-Šidák multiple comparisons adjustment as appropriate.

Discriminant analysis was employed for classification purposes. Feed-forward perceptron artificial neural networks were conducted with 200 iterations & 50 repeats per network. Support vector machines used (nonlinear) radial kernelized hyperplanes constructed from tune grids. Cross validation used repeated K-folds, LOOCV, and LGOCV/Monte Carlo methods. Missing data was assessed for MAR, MNAR, MCAR status and imputation approached by multiple imputation by chained equations (MICE). Ultimately, imputed datasets were not required during analysis.

## 3. Results

We used a suite of *ex vivo* assays to define the DDR landscape in ovarian cancer to see whether it could predict clinical outcomes.

Two separate patient cohorts were compiled. An “Optimisation cohort” was used to develop assays and explore any correlation between DDR and patient outcome. This was followed by an “Expanded cohort” which was used to carry out an extended analysis of the DDR and validate correlations seen with the Optimisation cohort.

### 3.1. Study pipeline and patient cohorts

The Optimisation cohort comprised 25 ovarian cancer patients recruited between 2011 and 2017. The Expanded cohort comprised 29 ovarian cancer patients recruited between 2018 and 2020. High-grade serous ovarian cancer (HGSOC) subsets were extracted from both cohorts and termed HGS-Optimisation and HGS-Expanded respectively to enable focused analyses (Figure 1).

**Figure 1.**
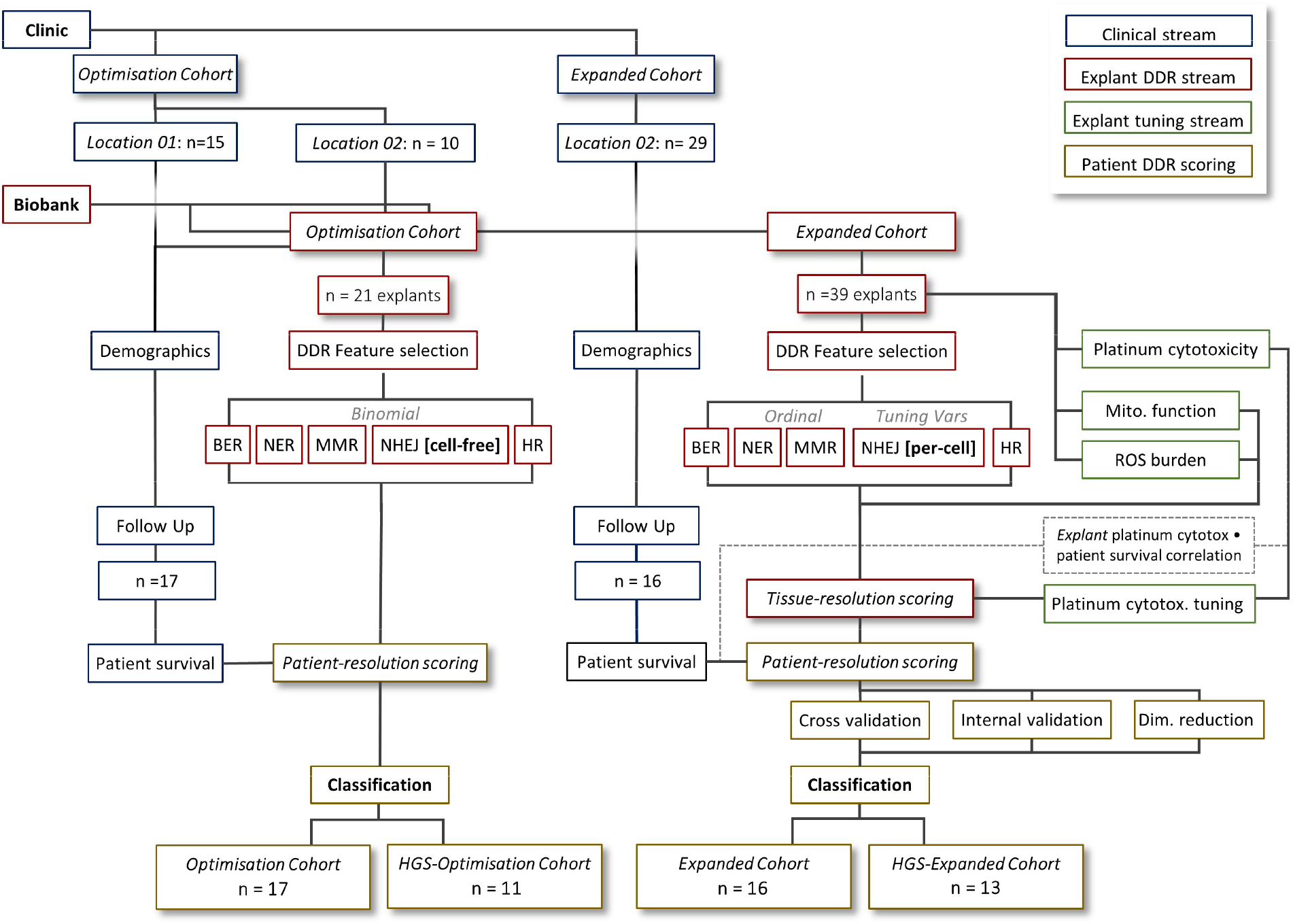
Research flow diagram. The pipeline followed for Optimisation and Expanded cohort research is described. Blue modules denote the clinical stream (see table 1). Maroon modules denote the biobank explant DDR stream. Green modules denote non-DDR explant experiments with classification tuning utility. Gold modules are final patient classification.

PFS and OS follow-up data was available for 17 and 11 patients in the Optimisation and HGS-Optimisation cohorts whilst data for 16 and 13 patients were available for the Expanded and HGS-Expanded cohorts respectively. With the exception of performance status, no significant cohort differences were observed for disease characteristics, physiological parameters and treatment pathways between cohorts (Table 1 and supplementary data section 2). All patients were treated with a combination of surgery and platinum based chemotherapy. Tissue samples were taken at the time of surgery with 24/54 (44%) of patients having primary surgery whilst the remainder had surgery after three cycles of neoadjuvant chemotherapy.

**Table 1.**
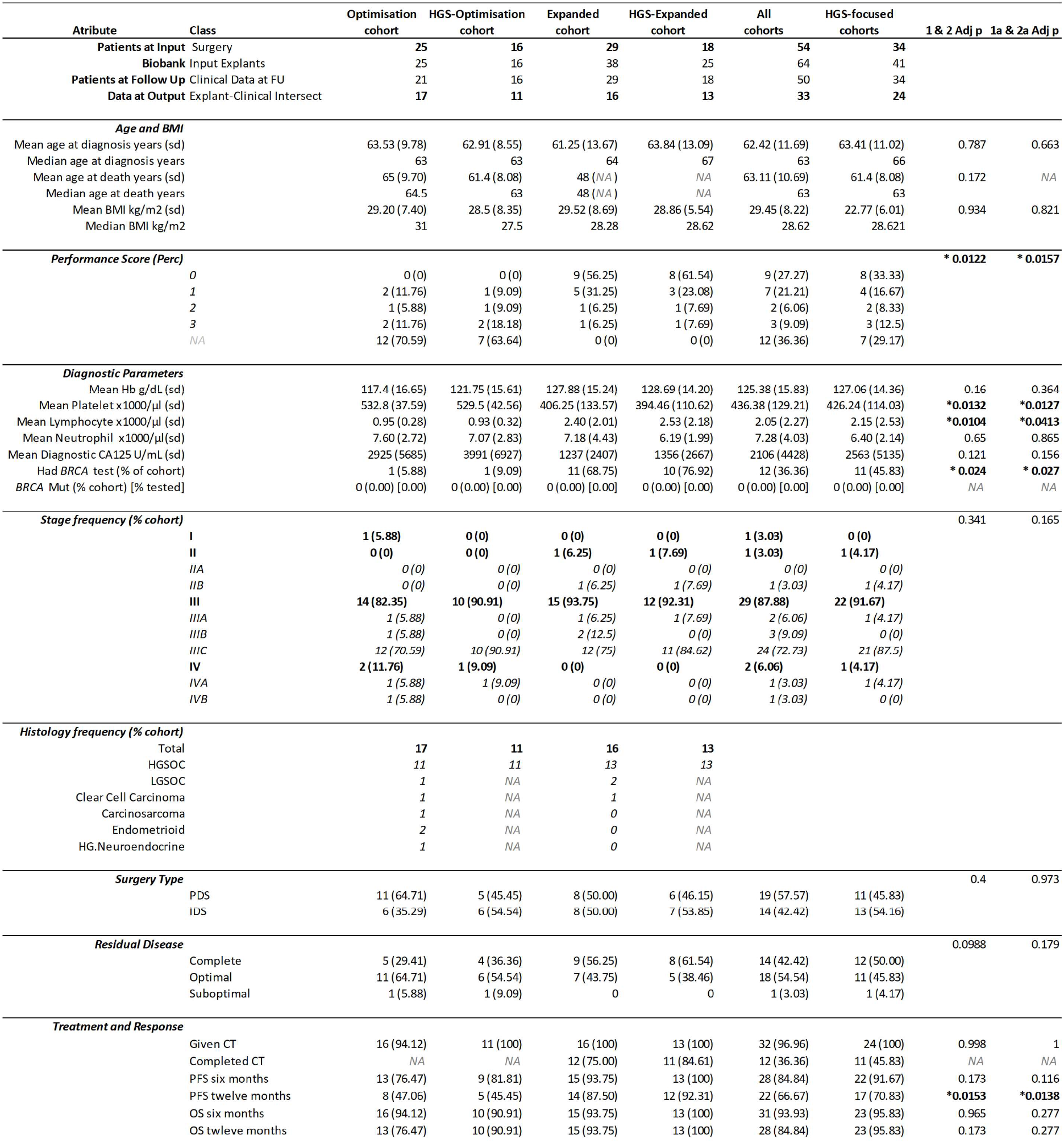
Patient cohort demographics. Clinical demographics for patients in this study. Optimisation and Expanded cohorts comprise any histology ovarian cancer. HGS-Optimisation and HGS-Expanded cohorts comprise HGSOC only.

### 3.2. Optimisation cohort and preliminary findings

For the Optimisation cohort, 25 explants were established and the *ex vivo* assay suite was used to generate binomial *competent* or *defective* scores for HR, NHEJ, BER, NER & MMR pathway states. Discriminant analysis used these DDR scores to classify overall survival of the 17 Optimisation cohort patients to achieve an AUC of 0.856 (supplementary data section 3). This performance improved when limited to the HGS-Optimisation cohort to achieve an AUC of 0.950, which was driven by a single misclassified sensitive patient. Further details are provided in supplementary data section 3 illustrating the importance of functional DDR pathway capacity assays.

### 3.3. Expanded cohort: a global DDR pathway capacity signature

In order to further explore and refine our DDR pathway capacity assays we developed an Expanded cohort pipeline, Figure 1, whereby 38 explants were established from 29 patients, (Table 1 and supplementary data section 4) of which 62% were HGSOC and the remaining 38% comprising other ovarian cancer histotypes (supplementary data section 4A & B). Our explant establishment success rate was 88% with 22 patients having a single explant, five patients with two explants, and two patients with three explants (supplementary data section 4C & D). Thus, in contrast to the Optimisation cohort this Expanded cohort allowed an exploration of tissue site heterogeneity. Details of anatomical site of origin of explants, and growth characteristics are given in supplementary data 4E &4F.

We profiled explant cultures using an extended functional DDR capacity assay panel which, in addition to the five DDR pathway assays, also included assays to determine explant platinum cytotoxicity state, explant response to ROS burden, and explant mitochondrial membrane potential, Figure 2. Details of the scoring system for each assay are outlined in methods and supplementary methods 1.5. Assays showed no preferential bias between explant tissue types or broad tissue types (supplementary data section 5A-5G) thus demonstrating technical robustness.

**Figure 2.**
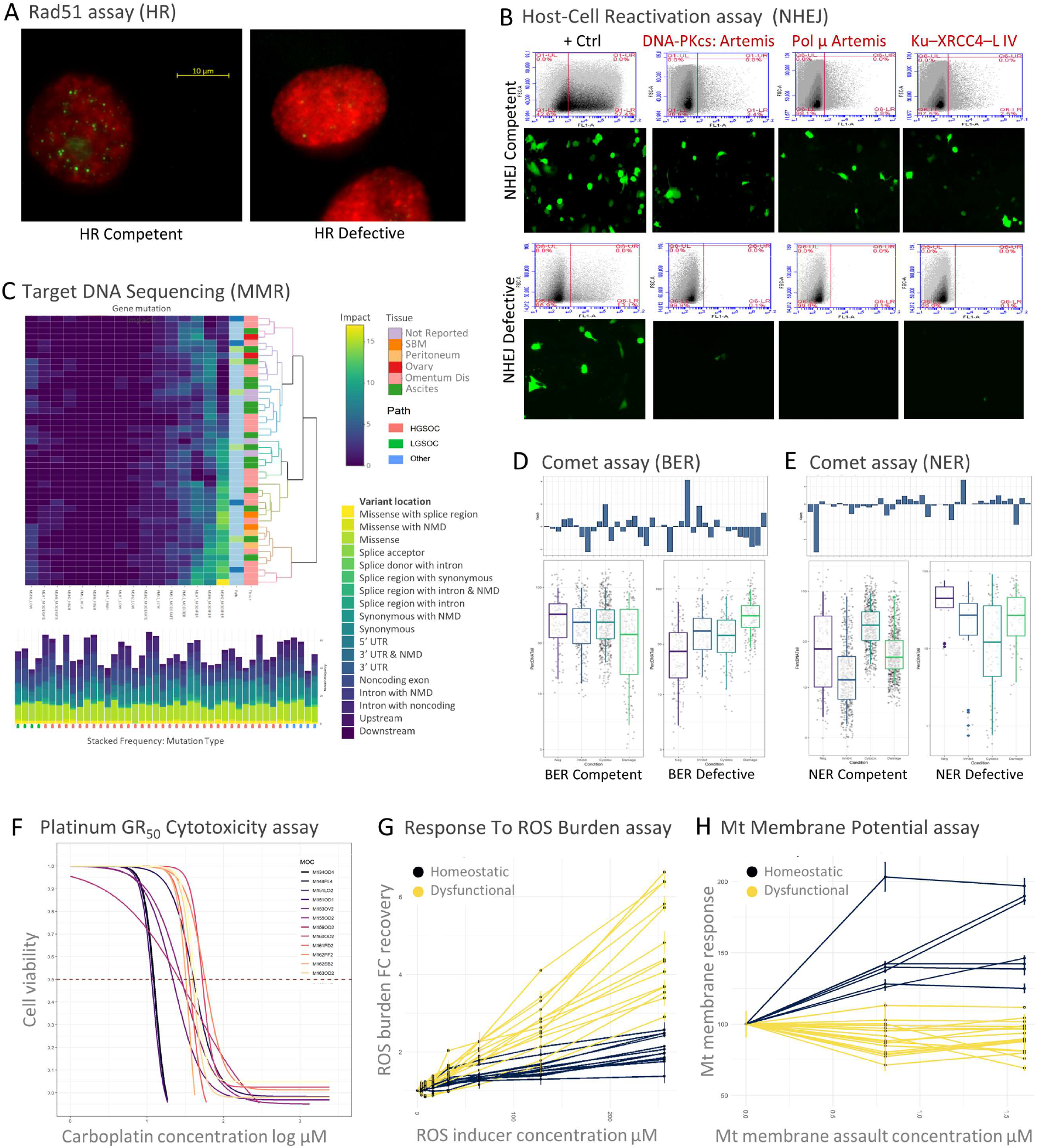
A:E: Representative explant-resolution DNA Damage Response capacities in ovarian cancer. F:H: Representative explant-resolution cytotoxic and metabolic tuning assays in ovarian cancer. **A**. Rad51 assay (HR). Immunofluorescence-based explant cell scoring for Rad51 localisation and nucleoprotein filament formation following DNA DSB. Extent of repair was used to score explant HR pathway capacity. **B**. Host-Cell Reactivation assay (NHEJ). Plasmid GFP expression cassettes were subject to three double-strand break conformation types and introduced into explant cells. Repaired plasmid double-strand breaks restores the cassette to permit GFP expression. Extent of repair was monitored qualitatively by fluorescence microscopy and quantitatively by flow cytometry. Each repaired conformation necessitates a particular cohort of NHEJ pathway components (maroon font). Extent of repair was used to score explant NHEJ pathway capacity. **C**. Target DNA sequencing (MMR). Mutation status of MMR cohort genes was extracted from exome sequencing and assessed by mutation type, location, and frequency. VEP and SnpEff Impact Scores were used to score explant MMR pathway capacity. **D & E**. Comet assays for BER and NER respectively. Explants were subjected to BER or NER recipient single-strand break DNA damage in presence or absence of pathway blockade. Extent of repair was measured by high-throughput comet assays (≈7000 comets typically per explant) and comet tail percent DNA metrics were used to score BER and NER pathway status. **F**. Explant platinum cytotoxicity GR_50_ plots [33]. A range of cell viability is evident across explant cultures following treatment with increasing concentrations of carboplatin. **G**. Explant response to reactive oxygen species (ROS) burden. Cells were treated with increasing concentration of the ROS inducer TBHP. Cells were scored as homeostatic or dysfunctional based upon DCFDA-derived extent of ROS retained in cells following a period of recovery. **H**. Explant mitochondrial membrane potential status. Mitochondrial (Mt) membrane activity following Mt assault (percent-normalised to negative control cells). Homeostatic explant Mt membranes responded to assault c.f. dysfunctional explants whose Mt membranes demonstrated no material response to membrane assault.

### 3.4. The DNA Damage Response pathway landscape in ovarian cancer explant cultures is varied

Each explant demonstrated at least one dysfunctional DSB pathway and one dysfunctional SSB pathway. HR defective status was associated with a greater dysfunction in the remaining pathways such that at least three DDR pathways per explant culture were highly perturbed when HR pathways are defective, Figure 3A.

**Figure 3.**
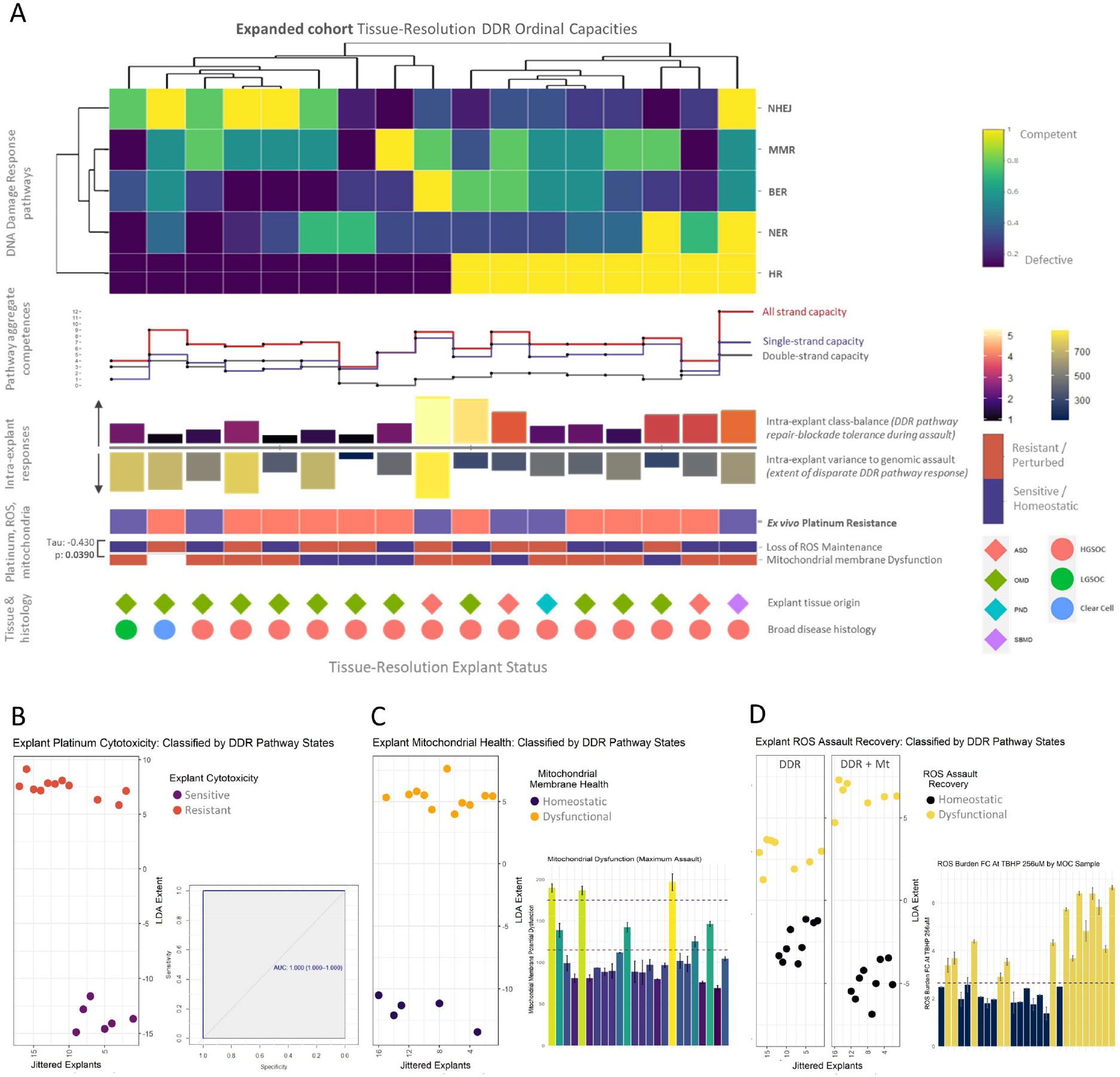
Expanded cohort tissue-resolution explant DDR capacities with associated cytotoxicity and metabolic classifications. **A**. DNA Damage Response (DDR) pathway states for Expanded cohort explants. Each column is an explant and each row is a DDR pathway capacity. A range of defective pathway signatures are evident. Clustering resolves Host-Cell Reactivation (NHEJ) and Rad51 (HR) assays to almost diametrically-opposed DSB repair capacity states. Intra-explant class balance bar plots depict extent-of-cell-viability survivorship following repair-blockade during genomic assault. **B**. Explant platinum cytotoxicity was completely classified by the five DDR pathway capacity states (discriminananalysis ROC AUC of 1). **C**. The five DDR pathway capacity states completely classify Mt membrane health (discriminant analysis ROC AUC of 1). **D**. Explant response to ROS assaults was described by the five DDR pathway capacity states (AUC of 1) with distinct but limited discrimination distances (left plot). A marked increased discriminatory distance was obtained when supplemented with Mt membrane status (right plot). An explant Mt membrane state and its ability to recover a ROS burden were inversely correlated (p 0.0390).

The extent-of-cell-viability survivorship following repair-blockade during genomic assault was examined using intra-explant class balance, Figure 3A. A wide range of survivorship was evident across the explants. HR competent explants demonstrated increased tolerance to repair-blockade during assault with higher overall cell survivorship in contrast to HR defective explants which contained an overall reduced cell survivorship bias. These observations are mechanistically in-line with HR competence states and associated tumour resistance.

Next, the cell population extent-of-variance following genomic assault was examined using intra explant variance bar plots, Figure 3A. A wide range of intra-explant cell variance was evident across the cohort. HR competent explants exhibited decreased variance in their response to genomic assault. This contrasted with HR defective explants which demonstrated an overall increased per-cell variance range to steady genomic assault. This observed greater intra-explant inter-cell heterogeneity is in concordance with the greater loss of DDR pathway capacities in HR-defective cells in contrast to the HR-competent explant reduced variance due to tighter rebuttal to aggregation of genomic instability during tumour evolution.

Of the explant cultures, 11 out of 17 (65%) were determined as platinum resistant (Gr_50_>= 48). This was equally split across both HR competent and HR defective explants indicative that platinum relation to the DDR landscape is complex and is not a direct response to any individual DDR pathway capacity.

The explant response to a ROS burden was wide ranging across the DDR pathway explant signatures. Five out of eight (62.5%) HR competent and four out of nine (44%) HR defective explants were homeostatic for ROS burden control, Figure 3A.

The Mt membrane potential signatures were largely dysfunctional across the explant cohort. Four out of five (80%) of platinum resistant HR competent and two out of three (66%) platinum sensitive HR competent explants exhibited perturbed Mt membrane potentials. Three out of six (50%) platinum resistant HR defective explants and three out of three (100%) platinum sensitive HR defective explants exhibited perturbed Mt membrane potentials indicative of no direct platinum associations.

We observed a modest significant inverse correlation between the explant’s mitochondrial membrane health and its ability to recover from a ROS assault (Kendall’s tau: -0.430, p = 0.0390). One out of 17 (5.9%) explants was homeostatic for both ROS and Mt membrane function, while three out of 17 (17.6%) explants were perturbed for both ROS and Mt membrane function. 13 out of 17 (76%) displayed inverse states, Figure 3A.

### 3.5. The ovarian cancer explant DDR pathway landscape classifies *ex vivo* platinum cytotoxicity and metabolic activity states

We next investigated whether the explant platinum cytotoxicity status could be predicted by the observed DDR pathway capacity signature. Using discriminant analysis, we discovered that explant platinum cytotoxicity was completely classified by the five DDR pathway capacity states determined by our *ex vivo* assays and achieved an AUC of 1, Figure 3B. The greatest DDR pathway-derived discriminant coefficients (supplementary data section 6A) were NHEJ status (11.42) and MMR status (4.34) indicative of the important involvement of both DSB and SSB DDR pathway status in cell tolerance of platinum agents. The greatest non-DDR-capacity discriminant coefficient was intra-explant extent-of-cell-viability survivorship (13.31) which mirrors an aggressive cell viability phenotype and supports the relevance of survivorship bias metrics within explant analysis.

We discovered that complete discrimination of mitochondrial membrane health was achieved by the five DDR pathway capacity states (AUC of 1) (Figure 3C). The discriminatory distance was driven by HR (17.44) and NHEJ (15.71) DDR pathway-derived coefficients and by intra-explant extent-of-cell-viability survivorship (e.g., 25.28) (supplementary data section 6B).

An explant’s ability to respond to a ROS assault was described by the five DDR pathway capacity states (AUC of 1) with a distinct and limited discrimination distance (Figure 3D). Given (i) our observed inverse correlation with explant mitochondrial dysfunction (above), and (ii) the mechanistic link between mitochondrial activity-derived ROS generation and a cell’s efficacy to control cytoplasmic ROS levels, we supplemented the five DDR pathway capacity signatures with the known Mt membrane status and obtained a markedly greater discriminatory distance driven by a negative Mt membrane dysfunction coefficient (−9.38) (supplementary data section 6C).

### 3.6. Patient summary analysis of explant DDR pathway, cytotoxicity, and metabolic states

Next we examined the data at patient level by developing a patient score which accommodated the presence of multiple explants per patient. An algorithm ensured consistent summarising and was weighted by DDR capacity values, range of intra-explant variances, inter-explant heterogeneity, and the degree of class-balance cell survivorship (supplementary methods 1.5).

Dendrogrammatic clustering of patients by DDR capacity states resolve the two DSB pathways to be diametrically opposed (Figure 4A), with at least 1 DSB and 1 SSB defective pathway for each patient. HR defective status associated with a greater dysfunction in the remaining pathways such that at least three DDR pathways per patient are highly perturbed when HR pathways are defective. No significant difference was observed between the extent of patient DDR pathway competencies and patient IDS or PS treatment routes (p value ranges: 0.308 to 0.874; supplementary data section 7) indicative that stand break repair competencies profiled here were tumour-evolution derived events rather than IDS CT-derived occurrences. No patient had complete dysfunction across the entire DDR pathway complement.

**Figure 4.**
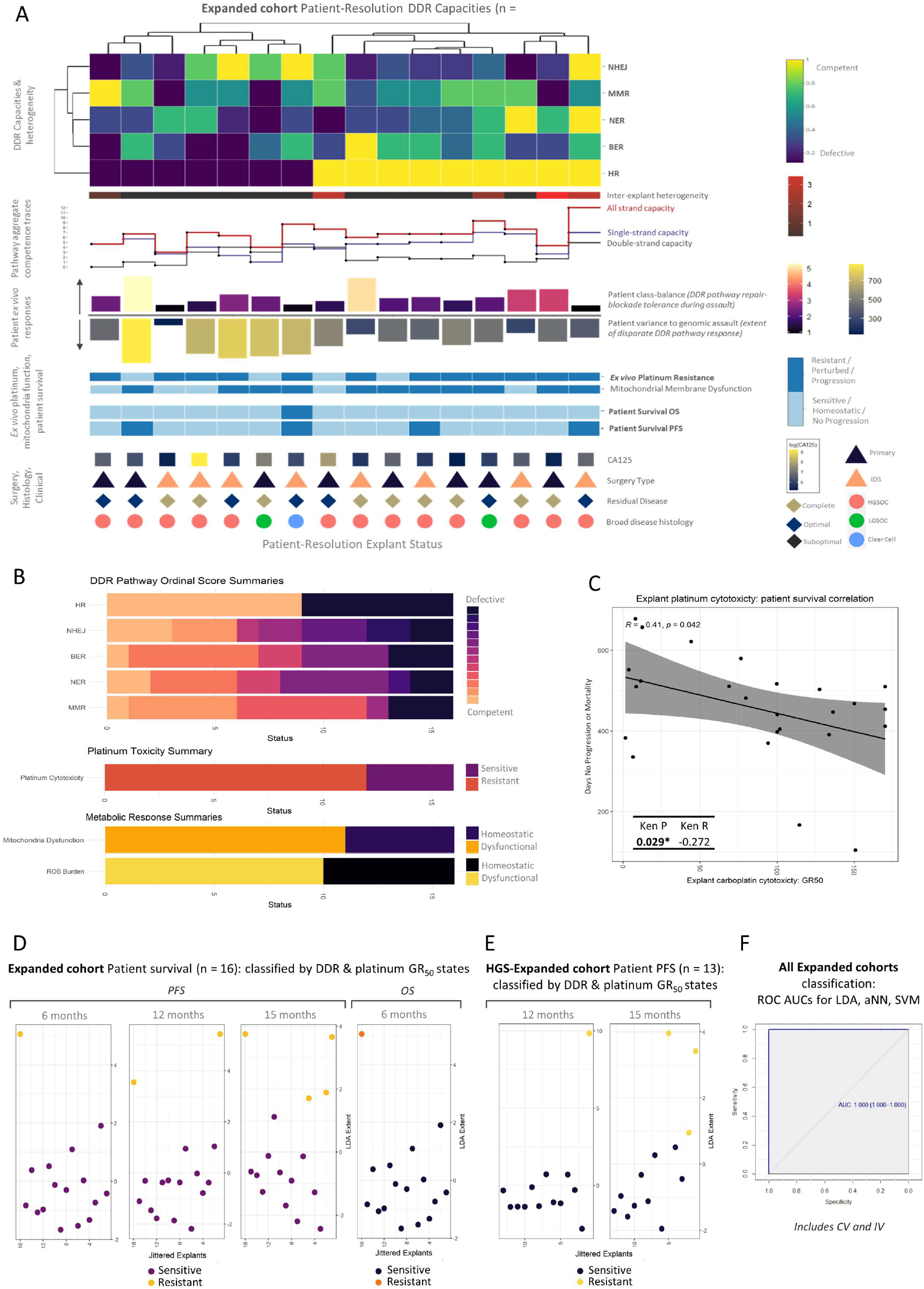
Expanded cohort patient-resolution explant DDR capacities with platinum survival correlation and patient classifications. **A**. DNA Damage Response landscape in relation to patient survival. Each column is an Expanded cohort patient and each row is a DNA damage response pathway capacity **B**. Summary bar plots of frequency of DDR pathway capacity metrics, platinum cytotoxicity, mitochondrial dysfunction, and response to ROS burden. **C**. A modest inverse correlation between explant platinum cytotoxicity and patient progression-free or overall survival was observed (r = 00.41, Pearson p = 0.042; Kendall’s tau p = 0.029). **D**. DDR pathway capacity signatures and platinum cytotoxicity states can completely classify Expanded cohort patient progression-free survival and overall survival status by discriminant analysis at six, 12, and 15 months (ROC AUC of 1 – see **F**). **E**. DDR pathway capacity signatures and platinum cytotoxicity states can completely classify HGS-Expanded cohort patient progression-free survival by discriminant analysis at 12 months and 15 months (ROC AUC of 1 – see **F**). **F**. Complete classifications were maintained for Expanded and HGS-Expanded cohorts up to 15-month progression-free survival with the use of nonlinear feedforward multilayer perceptron artificial neural networks (aNN) or radial basis function (RBF) kernel support vector machines (SVM) in conjunction with cross validation and internal validation (ROC AUC of 1). Expanded cohort six-month PFS and six-month OS, and HGS-Expanded cohort 12-month PFS each contain a single resistant patient and thus were unsuitable for data partitioned internal validation.

The inter-explant heterogeneity score denotes that four out of nine (44.4 %) HR competent patients demonstrated inter-explant heterogeneity in contrast to one out of seven (14.2 %) HR defective patients, Figure 4A. The inter-explant heterogeneity score did not directly associate with patient PFS or OS.

Wide ranges of repair-blockade tolerances during genomic assault and intra-explant cell variances were retained across the patient cohort. Patients presented similar trends to explant cultures wherein HR-competent patients had overall increased viability tolerances in contrast to HR defective patients. HR competent patients presented decreased per-cell variance ranges in response to genomic assault in contrast to HR defective patients, Figure 4A.

Follow up confirmed that 12 out of 16 (75%) patients were platinum resistant (relapsed with a treatment free interval of less than six months) and three out of four (75%) patients who progressed were platinum cytotoxicity resistant. Seven out of nine (77.8%) HR competent and five out of seven (71.4%) HR defective patients were platinum cytotoxicity resistant which consolidated the explant-resolution observation that platinum relation to the DDR landscape is complex and is not a direct response to any individual DDR pathway capacity.

Mitochondrial membrane functionality was perturbed in seven of nine (78%) HR competent patients but only four of seven (57%) in HR defective patients. Every patient who relapsed or died exhibited perturbed Mt membrane functionality, Figure 4A.

To summarise these findings, DDR pathway dysfunction was wide ranging across our patient cohort. Seven out of 16 (44%) patients were HR pathway defective while a range of NHEJ, BER, NER, and MMR competence scores were evident with most patients presenting a range of partially perturbed or basal competence signatures. The majority of the patient cohort presented aberrant ROS control and dysfunctional mitochondrial membrane potentials, Figure 4B.

We next assessed whether explant platinum cytotoxicity assays offer direct correlation utility with patient survival. A modest inverse correlation exists between explant platinum cytotoxicity and progression-free or overall survival (r = -0.41, Pearson p = 0.042; Kendall tau = -0.272, Kendall p = 0.029, Figure 4C).

### 3.7. Ovarian cancer patient PFS and OS can be classified by the DDR landscape and platinum cytotoxicity states

We next sought to determine whether patient-summarised DNA Damage Response pathway capacity signatures and associated platinum cytotoxicity could be used to classify patient progression-free survival or overall survival. Figure 4D depicts patient survival classification for the Expanded cohort by summarised explant DDR capacity signatures and platinum cytotoxicity at six months, 12 months, and 15 months (the maximum follow-up duration available at this time). One patient progressed within six months, two patients progressed within 12 months, and four patients progressed within 15 months. A single Expanded cohort patient died within six months during this 15-month period. DDR pathway capacity signatures and platinum cytotoxicity states can completely classify Expanded cohort patient progression-free survival and overall survival status by discriminant analysis (AUC of 1; Figure 4F). This observation remained true when limited to patients with HGSOC pathology. Figure 4E depicts HGSOC patient survival classification by summarised explant DDR capacity signatures and platinum cytotoxicity at 12 months and 15 months. No HGS-Expanded cohort patients progressed within six months, one patient progressed within 12 months, and three patients progressed within 15 months. No HGS-Expanded cohort patients died within this 15-month period. DDR pathway capacity signatures and platinum cytotoxicity states completely classified HGS-Expanded cohort patient progression-free survival by discriminant analysis (AUC of 1; Figure 4F).

To assess whether the observed classification efficacy was limited to an individual approach, we adopted additional non-linear machine learning approaches that included feedforward multilayer perceptron artificial neural networks (aNN) and radial basis function (RBF) kernel support vector machines (SVM) in conjunction with cross validation and internal validation. Complete classifications were achieved for Expanded and HGS-Expanded cohorts up to 15-month progression-free survival by aNN and RBF-SVM approaches (AUC of 1; Figure 4F).

### 3.8. Dimension reduction identifies DDR drivers of PFS

Principal component analysis (PCA) dimension reduction identified that HR, NHEJ, NER, and MMR capacities contribute to patient resistance classification, Figure 5. Non-DDR capacity metrics such as intra-explant extent-of-variance, survivorship class balance, and explant platinum cytotoxicity also contributed strongly.

**Figure 5.**
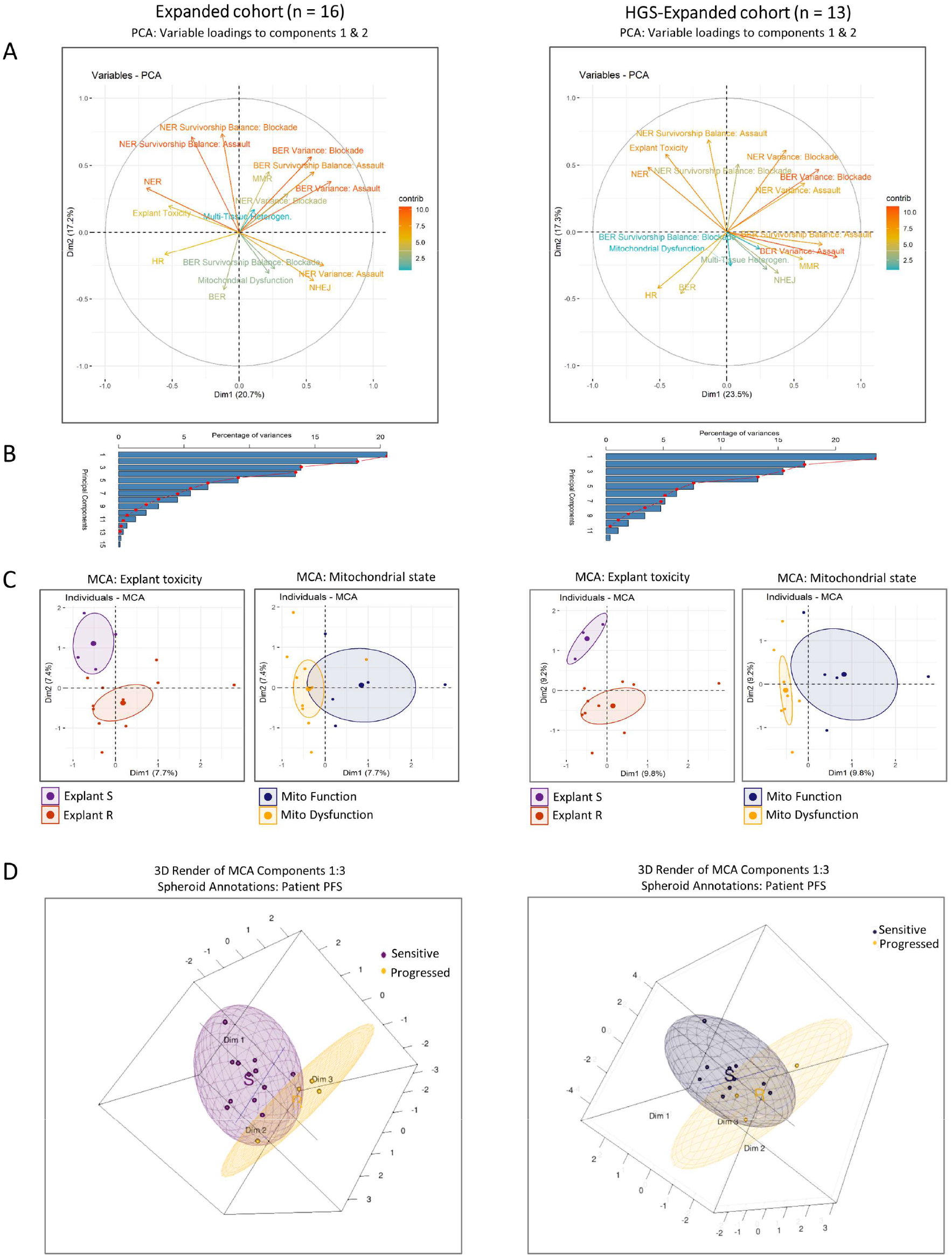
DDR capacity contributions to patient classification: Dimension Reduction. Left column: Expanded cohort. Right column: HGS-Expanded cohort. **A**. Principal component analysis (PCA) dimension reduction variable loading compass plots of DDR pathway capacity functional assay panel to patient resistance, coloured by their contributions to components one and two. HR, NHEJ, NER, and MMR capacity states occupy distinct directionally within both cohorts and contribute the most. Non-DDR capacity metrics (intra-explant extent-of-variance, survivorship class balance, and explant platinum cytotoxicity) also strongly contribute. **B**. Associated PCA variance scree plots. **C**. Multiple correspondence analysis (MCA) dimension reduction of platinum cytotoxicity and mitochondrial membrane dysfunction states classified by DDR pathway capacities. Platinum cytotoxicity ellipses (left insets) present clear separation in the Expanded and HGS-Expanded cohorts. Mitochondrial membrane dysfunction ellipses (right insets) present distinct but overlapping regions in the Expanded cohort; and fully partitioned ellipses in the HGS-Expanded cohort. **D**. Patient PFS classification by explant DDR and platinum status constructed from the first three MCA components and plotted into three-dimensional space. Spheroids provide patient survival classification group clusters. Distinct spheroids are visible for Expanded and HGS-Expanded cohort patient survival classes.

Finally, multiple correspondence analysis (MCA) dimension reduction was used to provide an unsupervised view of the data. MCA identified separating clusters corresponding to explant cytotoxicity and mitochondrial status, Figure 5C and progression free survival, Figure 5D thus confirming the ability of DDR signatures to predict *ex vivo* and clinical outcomes.

## 4. Discussion

We present here a suite of functional assays to describe the status of the five canonical DDR pathways. Combined with *ex vivo* platinum cytotoxicity these functional DDR capacity assays completely classified and predicted ovarian cancer patient progression-free or overall survival.

Chemosensitivity is a multifaceted phenomenon yet the intrinsic ability of a cell to repair DNA in response to chemotherapy-induced damage is a fundamental factor. Although extensively described, the role of homologous recombination has generally been studied in isolation and the role of alternative pathways are often overlooked. Here, we have attempted to employ a range of assays which reflect a holistic picture of the DDR at an explant and patient level.

For a disease that is hallmarked by extensive chromosome instability it is perhaps not surprising that discrete signature patterns of DDR dysregulation were not seen across the cohorts of high grade serous cancers studied here. Instead, almost all permutations of DDR dysregulation are seen, with a preponderance towards more pathway dysregulation in the HRD tumours than in the HRC cohort. Our finding that the two double strand repair pathways show near mutual exclusivity is interesting and suggests that strategies to identify druggable targets in the NHEJ pathway would have clinical utility in complementing PARP inhibitors, which preferentially target HRD tumours.

We also included ovarian tumours with a non-high-grade serous morphology. These tumours have different molecular drivers including kras/braf and PTEN [36] [37] [38]. That these tumours also display similar degrees of DDR dysregulation to HGSOC is encouraging that a DDR signature may be a useful overlay to morphology in a wide variety of solid tumours.

All tumours used in this study were collected during primary treatment and all patients received carboplatin combined with paclitaxel as the mainstay of their chemotherapy. At the time of study recruitment PARP inhibitor therapy was not available as maintenance therapy with no patient receiving this before first relapse. Our ability to classify and predict outcome for this cohort therefore represents a model for predicting response to platinum/taxane combination therapy only. Nevertheless, this model shows promise as a clinical tool to predict chemosensitivity to these agents, both of which are used extensively in the relapse setting.

The models generated here compare favourably with other methods of classification that have been proposed for high grade serous cancer including DNA [21, 39] and RNA [40] based models. Moreover the functional approach to these assays is justified by the incremental improvement in prediction seen with these assays compared to recent studies using an IHC approach [41], and similar results in glioblastoma multiforme [42] with a panel including the host-cell reactivation system used here [25].

It is unsurprising that multiple pathways are required to generate an accurate predictive signature. Although the association of HR repair status with chemoresistance is known, an emphasis on any single pathway is mechanistically insufficient to capture compensatory and reciprocal activity across the remaining DDR pathways [5]. On the one hand XRCC1 actuation within the predominant SSB repair PARP-dependent BER pathway associates with poor clinical outcomes, whilst XRCC1 deficient [43] or APE1/Ref-1 silenced cells [44] are sensitized to cisplatin. Similarly, loss [45] or alternative-splicing [46] of the NER pathway component ERCC1 or loss of XPF [47] associates with platinum sensitivity in cells. These SSB pathway reports appear to align with the HR pathway notion that functional repair associates with poorer outcomes. Conversely, MMR has been reported to both offer [8] and not offer [48] prognostic significance, whilst NHEJ defectiveness can associate with sensitive and resistant phenotypes which can occur through 53BP1-derived NHEJ loss to reactivate HR competence via a *BRCA*-independent manner [49].

The association and relevance of tumour heterogeneity to platinum chemoresistance has come to prominence in recent years. Patch *et al* [50] evaluated the whole-genome landscape of chemoresistance ovarianchemoresistant ovarian cancer and reported relatively low point mutations amongst extensive genomic rearrangement, high ORF breakage and multiple independent *BRCA* reversion mutations within the same patient’s tumour. Extensive heterogeneity exists across different metastases within the same patient [51] and across individual tumour architecture which likely precludes platinum-instigated immune activation in the tumour microenvironment and contributes to chemoresistance [52]. We report here that HR defective *ex vivo* cultures exhibit increased intra-tumour variance indicative of a highly heterogeneous phenotype wherein greater instability is likely driven by loss of repair. Conversely HR competent explants displayed greater uniform states with increased tolerance to repair-blockade during assault indicative of resistance potential. Furthermore, platinum cytotoxicity responses were predominantly driven by NHEJ & MMR competence scores and class balance survivorship. This implicates the importance of both DSB and SSB repair pathways to the resistant phenotype and illustrates the benefit of monitoring cell survivorship in assays.

Mitochondria are central to a host of fundamental physiological processes and contribute to ROS generation and control [10]. Increased ROS abundance & oxidative stress are frequent events in ovarian cancer, and chemotherapy elevates ROS levels to alter cancer cell redox-homeostasis [12]. We observed that 78% of HR competent patients harboured dysfunctional Mt membrane in contrast to 57% of HR defective patients. Within the platinum cytotoxic resistant subset this partition further increased wherein 86% of HR competent patients were Mt membrane dysfunctional vs 40% in HR defective patients, and every relapse or mortality event contained perturbed Mt membrane functionality. Furthermore, membrane action and ROS assault recovery were significantly inversely correlated. Of interest, ovarian cancer *ex vivo* mitochondrial membrane health can be fully classified by the DDR landscape signature which was driven predominantly by HR, NHEJ, MMR, and intratumour class-balance. This was surprising but could be explained by the possibility that, in the presence of a DDR competent phenotype, a membrane-perturbed oncogenic mitochondrion will drive a loss of ROS cytoplasmic homeostasis and thus evolutionarily-enrich for ROS tolerances beyond the necessary chemo-induced cytotoxic threshold to promote chemoresistance.

Together, our findings show that the complexity and heterogeneity of the DDR response in cancer can be untangled using a suite of assays representing both the five major pathways and more global representations of the DDR. These can be combined to provide accurate predictions of response to standard chemotherapy regimens and so suggests clinical utility, particularly at clinical decision points, such as for relapsed disease, where the decision to treat is not always obvious. Although we have limited this study to ovarian cancer, the inclusion of multiple subtypes with different molecular drivers, suggests that this work will also be applicable to other cancer types.

