## supplementary methods for "The heterogeneity of the DNA damage response landscape determines patient outcomes in ovarian cancer"

### 1. Methods

Supplementary methods referenced from manuscript methods section are detailed here.

#### 1.1. Explant characterisation antibodies

| Antibody target | Species | Company | Cat. number |
| --- | --- | --- | --- |
| Anti-Ca125 | Mouse monoclonal | Abcam | ab1107 |
| Anti-Pax8 | Mouse monoclonal | Abcam | ab92547 |
| Anti-Vimentin | Rabbit Monoclonal | Abcam | ab92547 |
| Anti-Pan Cytokeratin | FITC-conjugated Mouse Monoclonal | Merck | cbl234f |
| Anti-Mouse IgG | Goat, Alexa Fluor 546 | Invitrogen | A-11003 |
| Anti-Rabbit IgG | Goat, Alexa fluor 488 | Invitrogen | A-11008 |

#### 1.2. Rad 51 assay (HR) antibodies

| Antibody target | Species | Company | Cat. number |
| --- | --- | --- | --- |
| Anti-Mouse IgG | Goat, Alexa Fluor 546 | Invitrogen | A-11003 |
| Anti-Rabbit IgG | Goat, Alexa fluor 488 | Invitrogen | A-11008 |

#### 1.3. *In vitro* cell extract assay (NHEJ) (cohort 1 & 1a)

Refer to ZF’s SOPs.

#### 1.4. Host-cell reactivation assay (NHEJ) preparations (cohort 2 & 2a)

The Host-cell Reactivation system (Nagel *et al* [1]) was further expanded to provide per-cell quantitative NHEJ pathway capacity monitoring of blunt end, 5’-3’ and 3’-5’ discontiguous de-phosphorylated mismatched overhang DNA double-strand breaks.

###### Host cell plasmid preparation

XL1-Blue supercompetent cells (200236; Agilent) were transformed with reconstituted pCMV6-AC-GFP plasmid (ps100010; Origene). Transformed cells were plated on LB agar (11508926; Invitrogen) containing 100µg/ml carbenicillin (C1389; Sigma) and incubated for 16 hours at 37°C. 20ml LB broth: 100µg/ml carbenicillin flasks were inoculated with transformed colonies and cultured for 8hrs at 37°C with 220rpm shaking. 1000ml step-up volumes were seeded from these and cultured under identical conditions for 16 hours in the presence of negative control flasks (non-transformed XL1-Blue cells, and no-cell media flasks). Plasmids were purified using endotoxin-free maxiprep kits and resuspended in endotoxin-free TE Buffer (12362; Qiagen). Purities and yields were assessed by UV spectrophotometry (NanoDrop ND-2000) and agarose gel analysis. ≈1000µg yields were obtained and plasmids were adjusted to 1000ng/ml concentrations.

EcoRI-HF (R3101S), SacI-HF (R3156S), KpnI-HF (R3142S), PmeI (R0560S), and Quick CIP (M0525) (New England Biolabs) were used with NEB CutSmart buffer and nuclease-free water to construct the three dephosphorylated double strand break (dsb) plasmid conformations provided below (condition 4 is uncut positive control plasmid).

| Condition | N-5’ | N-3’ | C-5’ | C-3’ | Blunt | Provides | Functioning NHEJ mediators likely required |
| --- | --- | --- | --- | --- | --- | --- | --- |
| 1 | EcoRI |  |  | KpnI |  | 5’ hang: 3’ hang | DNA-PKcs: Artemis endonuclease activity |
| 2 |  | SacI | EcoRI |  |  | 3’ hang: 5’ hang | Pol µ activity [Artemis likely required] |
| 3 |  |  |  |  | PmeI | Blunt | Ku–XRCC4–DNA ligase IV activity [Artemis not required] |

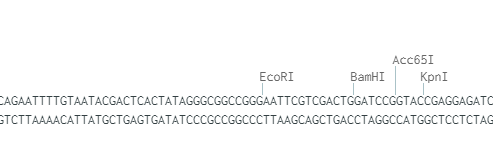

*15nt removed*

**Condition 1:** 5’ : 3’ double strand break

**Condition 2:** 3’ : 5’ double strand break

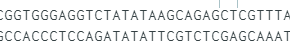

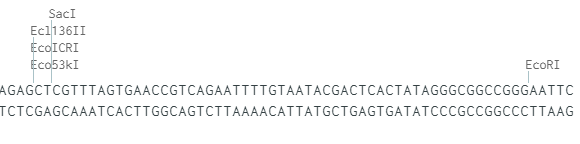

*52nt removed*

**Condition 3:** Blunt double strand break

*0nt removed*

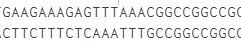

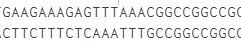

1µg 50µl restriction digest reactions were prepared with 20 units of each enzyme and 2n the suggested Quick CIP volume to obtain dephosphorylation. Reactions were incubated at 37°C for 4n the required time to achieve complete digestion and no more than 0.25n the minimum reported time prior to star activity, followed by heat-inactivation. Volumes were scaled to aliquots containing 10µg of plasmid as necessary in order to meet required yields.

2µg digested plasmid aliquots were resolved on 1% agarose gels alongside uncut reference plasmid and visualised by 1:1000 SYBR Safe (Thermo Fisher). dsb plasmid bands were gel extracted, purified, (QIAQuick Gel Extraction Kit 28704; Qiagen) and reconstituted in Endo-Free TE Buffer (12362; Qiagen). Plasmid purity and yields were assessed by agarose gel and spectrophotometry. From restriction digest preparation to digest purification (which incorporates gel-extraction), the inter-batch yield variation was 1% - 3% for the same plasmid conformation digests, and 1.7% - 7% yield variation between the three different plasmid conformations. Between-plasmid conformation purity ratio differences ranged between 0% - 0.6% (A260/280) and 1% - 3.5% (A260/230). These were indicative of robust and concordant dsb plasmid preparation and unlikely to contribute any bias towards final NHEJ quantitation analysis.

###### Explant plasmid transfection

Explant plasmid transfections were prepared with lipofectamine LTX Plus or 3000 (15338-100, 11668-027; Thermo Fisher) using Opti-Mem (11058021; Thermo Fisher) as per manufacturer’s guidelines. Conditions were determined empirically and used 52µl reactions with a final ratio of 2µl:0.5µg for LTX:DNA and 1.5µl:0.5µg for 3000:DNA. Replicate wells of 24 well plates were transfected with one of the four conditions (Uncut plasmid positive control, 5’-3’, 3’-5’, blunt-blunt) once explant cells were 85% confluent. Cultures were monitored at 24, 48, and 72 hours prior to analysis. Primary cells demonstrate documented challenges and a wide-range of transfection efficiencies via liposome-based transfection delivery methods. Any primary culture that exhibited suboptimal lipofectamine LTX transfection efficiencies were reanalysed using lipofectamine 3000 assays. Ultimately no explant was unamenable to optimal transfection assays.

###### Plasmid repair monitoring

Fluorescence-governed NHEJ pathway activity of each sample was assessed qualitatively by microscopy and quantitatively by flow cytometry. Cells were washed with PBS and visualised using a Zeiss Axio Observer Z1 microscope with Zeiss Zen 2.3 software. Parent cells confirmed absence of non-specific fluorescence whilst uncut plasmid provided positive transfection controls. For quantitative assessment the same samples were trypsinised, recovered, washed in PBS, and resuspended in 140µl 4°C sterile-filtered flow buffer (2% BS PBS). Samples remained on ice and cytometric fluorescence was determined on a BD Accuri C6 (Beckman Coulter). Cell-free control runs of media, PBS, flow buffer, deionised water, sheath fluid, decontamination fluid, cleaning fluid confirmed that zero non-specific events occurred at a frequency within two orders of magnitude below parent cell detection rate. The fluorescent detection extent of any non-specific events did not obfuscate negative control signals. Forward and side scatter gates were calibrated on parent cells and applied across corresponding explant NHEJ repair groups. Regions were calibrated on parent cells to determine their intrinsic fluorescence extent and regions were applied across corresponding NHEJ repair groups. Positive control samples confirmed explant transfection and permitted transfection efficiencies to be monitored. Fluorescence detection within dsb plasmid groups represents NHEJ pathway-governed processing and repair of DNA dsb to restore the GFP ORF and permit mRNA transcription and GFP protein expression. Extent of NHEJ pathway-governed repair was assessed with respect to both change in average fluorescence and change in number of cell events.

###### Average fluorescence intensity normalisation

Fluorescence change is concordant with traditional cell-free extract NHEJ assays and provides per-explant NHEJ pathway capacities. Average left-side region fluorescence intensities (defined as innate cell fluorescence as determined by parent cells) was subtracted from right-side region fluorescence intensity of dsb repair plasmids to provide between-explant normalised extent of fluorescence increase generated by NHEJ pathway-governed plasmid repair. Values were standardised to parent negative and transfection positive control conditions to provide percent-scaled capacities.

###### Cell event normalisation

Cell event monitoring enables per-cell resolution of intra-explant NHEJ repair capacity variance and thus enables detection of possible basal level NHEJ activity which would traditionally be obfuscated below aggregate-assay sensitivity thresholds. The ratio of event frequency of positive threshold cells to all sample cells was calculated to provide the proportion of cells within any sample that demonstrated repair activity. The event ratio of positive control samples represents the maximum achieved transfection efficiency per explant and was used to percentage-calibrate target condition group ratios.

###### NHEJ pathway competence score generation

An integrated NHEJ capacity scoring system was derived from equal-weighted contributions of cell event and fluorescent intensity standardised percent values. *Ordinal plasmid condition scores* were: Fully Defective (<5%); Basal Activity (≥5% & <15%); Low Activity (≥15% & <25%); Competent Activity (≥25% & < 35%); and High Activity (≥35%). *Plasmid condition scores* were combined to provide *Explant capacity scores* which measure overall explant NHEJ repair capacity. *Ordinal explant capacity scores* provided a 12-point scale (driven from the three input five-point scales).

#### 1.5. Statistical analysis, capacity scoring, classification, modelling, validation, dimension reduction

###### Software & hardware

R [2] packages used in addition to Base R are provided below.

| Row | Package | Citation |
| --- | --- | --- |
| 1 | tidyverse | [3] |
| 2 | splitstackshape | [4] |
| 3 | gridGraphics | [5] |
| 4 | gridExtra | [6] |
| 5 | gplots | [7] |
| 6 | ggpubr | [8] |
| 7 | viridis | [9] |
| 8 | RColorBrewer | [10] |
| 9 | plotly | [11] |
| 10 | webshot | [12] |
| 11 | htmlwidgets | [13] |
| 12 | ic50 | [14] |
| 13 | drc | [15] |
| 14 | PharmacoGx | [16] |
| 15 | GRmetrics | [17] |
| 16 | plotROC | [18] |
| 17 | pROC | [19] |
| 18 | MASS | [20] |
| 19 | nnet | [20] |
| 20 | car | [21] |
| 21 | caret | [22] |
| 22 | e1071 | [23] |
| 23 | Mice | [24] |
| 24 | VIM | [25] |
| 25 | vcd | [26] |
| 26 | rstatix | [27] |
| 27 | Hmisc | [28] |
| 28 | heatmaply | [29] |
| 29 | psych | [30] |
| 30 | rmcorr | [31] |
| 31 | corrplot | [32] |
| 32 | rcompanion | [33] |
| 33 | survival | [34] |
| 34 | survminer | [35] |
| 35 | PerformanceAnalytics | [36] |

###### Platinum cytotoxicity scores

Explant response to carboplatin was dichotomised using a GR_50_ value of <48µM for sensitive classification and ≥48µM for resistant classification. This value determined with reference to 1). The distribution of observed GR_50_ values across the explant cohort; 2). The range of carboplatin and cisplatin explant and cell line cytotoxicity values in the literature with consideration to their differing efficacies; 3). The typically observed increase in GR_50_ µM value vs the equivalent IC_50_ value per explant due to growth-rate correction within the model.

###### ROS scores

TBHP-negative DCFDA-treated replicate well Ex/Em 485/535 nm plate reader fluorescent values were subtracted from all TBHP treatment groups and ROS burden following increased TBHP treatment was converted to fold change events. Explants that were unable to recover the ROS assault to less than two-fold endogenous values were scored as perturbed.

###### Mitochondrial membrane health scores

Fluorescence values at Ex/Em 490/525 nm (loss of membrane potential) and 540/590 nm (active membrane potential) were used to classify mitochondrial membrane status. Cell-free background values were subtracted from all replicates and the intra-plate and inter-plate replicate mean and standard deviation assessed per parent or each drug concentration. Explant baseline mitochondrial membrane activity ratios and parent-normalised activity changes in response to the presence of drug were calculated and used to score mitochondrial status. Changes of less than parent-normalised 1.2FC across all drug treatments in conjunction with high parent cell mitochondrial membrane ratios were considered dysfunctional.

###### Patient summarised multi-explant culture scores

Patient-summarised multi-explant scores were generated by an algorithmic approach to accommodate potentially discordant explant capacities. BER and NER quantitative capacity score values were weighted by -Log KW p values and multi-explant average scores were generated. Binomial “has heterogeneity” and ordinal “consensus capacity” scores were introduced. MMR scores were averaged with an additional binomial “has heterogeneity” score and weighted for functioning explants. Averaged NHEJ ordinal scores were weighted for explant competence and binomial “has heterogeneity” and ordinal “consensus capacity” scores were introduced. Discordant binomial HR capacity scores were marked as “has heterogeneity” as required and a binomial “consensus capacity” score introduced for any odd-number-explant collapse. Intra-tissue assay variance was retained with weightings for greatest input variance values. Class-balance values were retained with weightings for greatest survivorship bias. Cytotoxicity values were weighted for resistance and binomial “has heterogeneity” and ordinal consensus scores introduced. ROS and Mt membrane scores were weighted for dysfunction alongside “has heterogeneity” and binomial “consensus capacity” scores.

23.	David Meyer, E.D., Kurt Hornik, Andreas Weingessel and Friedrich Leisch, *e1071: Misc Functions of the Department of Statistics, Probability Theory Group (Formerly: E1071), TU Wien.* 2020.

24.	van Buuren, S. and K. Groothuis-Oudshoorn, *mice: Multivariate Imputation by Chained Equations in R.* 2011, 2011. **45**(3): p. 67.

25.	Kowarik, A. and M. Templ, *Imputation with the R Package VIM.* 2016, 2016. **74**(7): p. 16.

26.	Meyer, D., A. Zeileis, and K. Hornik, *The Strucplot Framework: Visualizing Multi-way Contingency Tables with vcd.* 2006, 2006. **17**(3): p. 48.

27.	Kassambara, A., *rstatix: Pipe-Friendly Framework for Basic Statistical Tests.* 2021.

28.	Harrell.Jr, F.E., *Hmisc: Harrell Miscellaneous.* 2021.

29.	Galili, T., et al., *heatmaply: an R package for creating interactive cluster heatmaps for online publishing.* Bioinformatics, 2017. **34**(9): p. 1600-1602.

30.	Revelle, W., *psych: Procedures for Psychological, Psychometric, and Personality Research.* 2020.

31.	Bakdash, J.Z. and L.R. Marusich, *Repeated Measures Correlation.* Frontiers in Psychology, 2017. **8**(456).

32.	Wei, T. and V. Simko, *R package "corrplot": Visualization of a Correlation.* 2017.

33.	Mangiafico, S., *rcompanion: Functions to Support Extension Education Program Evaluation.* 2021.

34.	Therneau, T., *A Package for Survival Analysis in R.* 2020.

35.	Kassambara, A., M. Kosinski, and P. Biecek, *survminer: Drawing Survival Curves using 'ggplot2'.* 2020.

36.	Peterson, B.G. and P. Carl, *PerformanceAnalytics: Econometric Tools for Performance and Risk Analysis.* 2020.
