## supplementary data for "The heterogeneity of the DNA damage response landscape determines patient outcomes in ovarian cancer"

### Slide 1
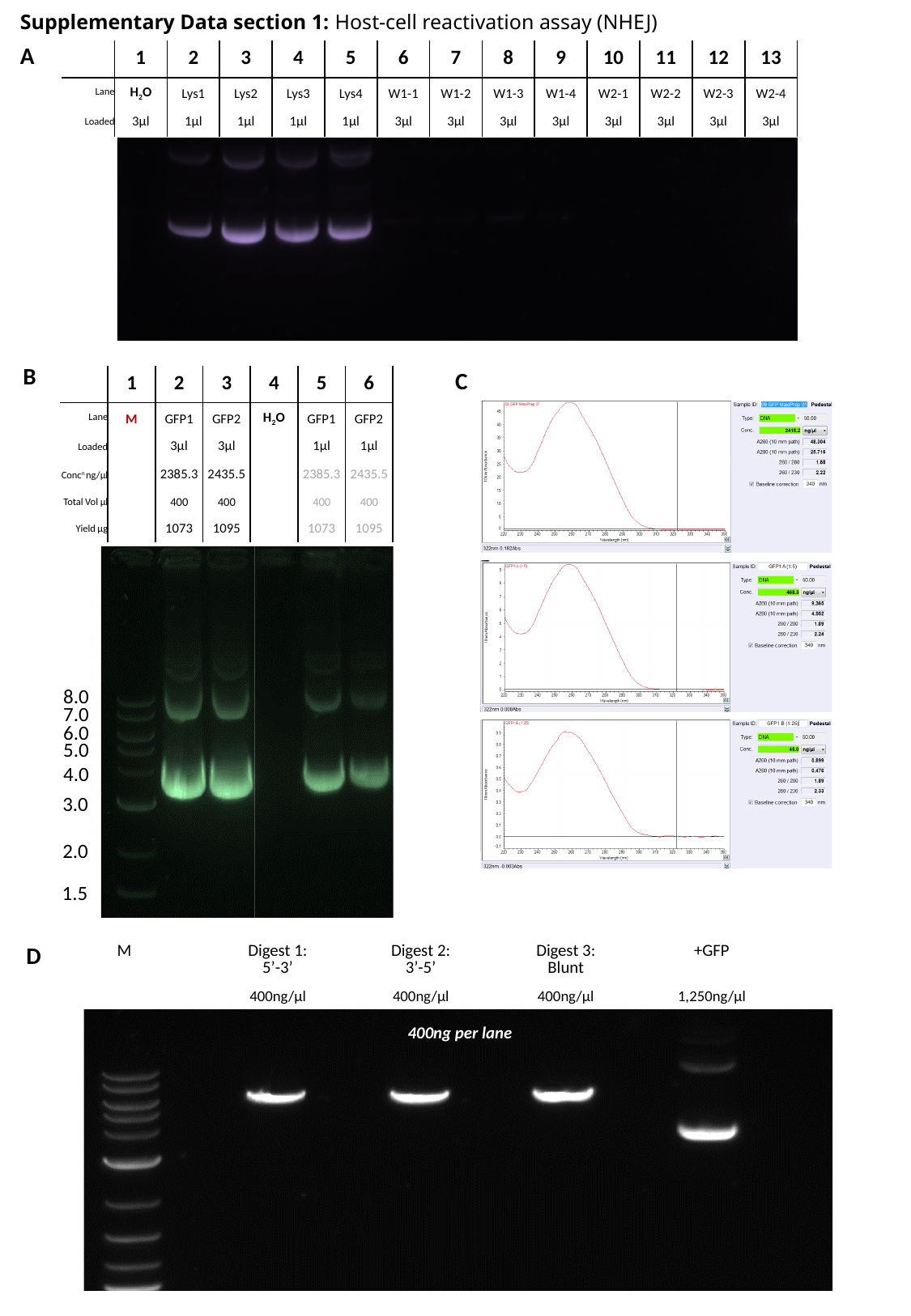

Supplementary Data section 1: Host-cell reactivation assay (NHEJ)
A
| | 1 | 2 | 3 | 4 | 5 | 6 | 7 | 8 | 9 | 10 | 11 | 12 | 13 |
| --- | --- | --- | --- | --- | --- | --- | --- | --- | --- | --- | --- | --- | --- |
| Lane | H_2_O | Lys1 | Lys2 | Lys3 | Lys4 | W1-1 | W1-2 | W1-3 | W1-4 | W2-1 | W2-2 | W2-3 | W2-4 |
| Loaded | 3µl | 1µl | 1µl | 1µl | 1µl | 3µl | 3µl | 3µl | 3µl | 3µl | 3µl | 3µl | 3µl |
B
C
| | 1 | 2 | 3 | 4 | 5 | 6 |
| --- | --- | --- | --- | --- | --- | --- |
| Lane | M | GFP1 | GFP2 | H_2_O | GFP1 | GFP2 |
| Loaded | | 3µl | 3µl | | 1µl | 1µl |
| Conc^n^ ng/µl | | 2385.3 | 2435.5 | | 2385.3 | 2435.5 |
| Total Vol µl | | 400 | 400 | | 400 | 400 |
| Yield µg | | 1073 | 1095 | | 1073 | 1095 |
8.0
7.0
6.0
5.0
4.0
3.0
2.0
1.5
D
| M | | Digest 1: 5’-3’ | | Digest 2: 3’-5’ | | Digest 3: Blunt | | +GFP |
| --- | --- | --- | --- | --- | --- | --- | --- | --- |
| | | 400ng/µl | | 400ng/µl | | 400ng/µl | | 1,250ng/µl |
| 400ng per lane | | | | | | | | |

### Slide 2
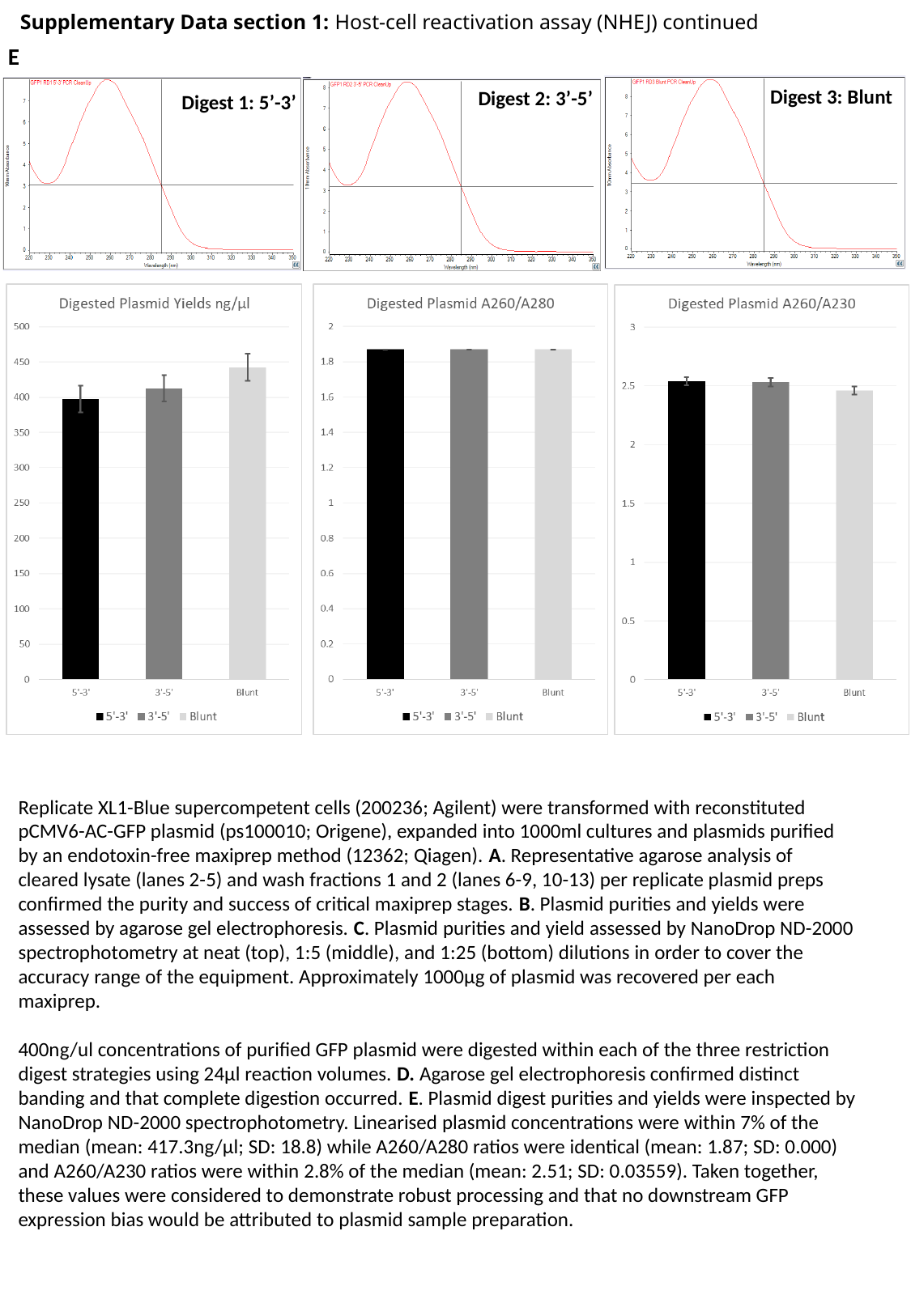

Supplementary Data section 1: Host-cell reactivation assay (NHEJ) continued
E
Digest 3: Blunt
Digest 1: 5’-3’
Digest 2: 3’-5’
Replicate XL1-Blue supercompetent cells (200236; Agilent) were transformed with reconstituted pCMV6-AC-GFP plasmid (ps100010; Origene), expanded into 1000ml cultures and plasmids purified by an endotoxin-free maxiprep method (12362; Qiagen). A. Representative agarose analysis of cleared lysate (lanes 2-5) and wash fractions 1 and 2 (lanes 6-9, 10-13) per replicate plasmid preps confirmed the purity and success of critical maxiprep stages. B. Plasmid purities and yields were assessed by agarose gel electrophoresis. C. Plasmid purities and yield assessed by NanoDrop ND-2000 spectrophotometry at neat (top), 1:5 (middle), and 1:25 (bottom) dilutions in order to cover the accuracy range of the equipment. Approximately 1000µg of plasmid was recovered per each maxiprep.
400ng/ul concentrations of purified GFP plasmid were digested within each of the three restriction digest strategies using 24µl reaction volumes. D. Agarose gel electrophoresis confirmed distinct banding and that complete digestion occurred. E. Plasmid digest purities and yields were inspected by NanoDrop ND-2000 spectrophotometry. Linearised plasmid concentrations were within 7% of the median (mean: 417.3ng/µl; SD: 18.8) while A260/A280 ratios were identical (mean: 1.87; SD: 0.000)
and A260/A230 ratios were within 2.8% of the median (mean: 2.51; SD: 0.03559). Taken together, these values were considered to demonstrate robust processing and that no downstream GFP expression bias would be attributed to plasmid sample preparation.

### Slide 3
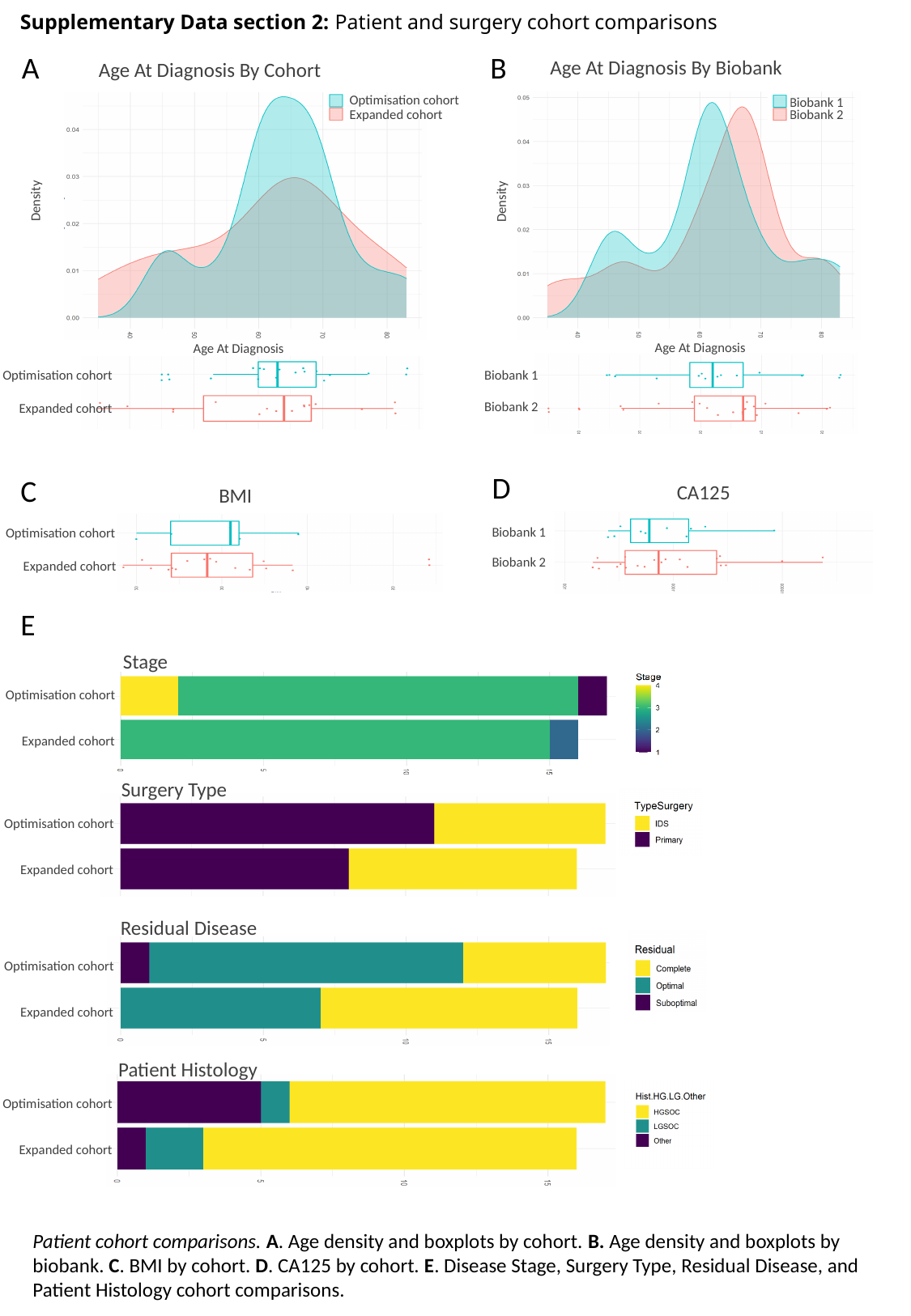

Supplementary Data section 2: Patient and surgery cohort comparisons
B
A
Age At Diagnosis By Biobank
Age At Diagnosis By Cohort
Optimisation cohort
Expanded cohort
Biobank 1
Biobank 2
Density
Density
Age At Diagnosis
Age At Diagnosis
Optimisation cohort
Expanded cohort
Biobank 1
Biobank 2
D
C
CA125
BMI
Biobank 1
Biobank 2
Optimisation cohort
Expanded cohort
E
Stage
Optimisation cohort
Expanded cohort
Surgery Type
Optimisation cohort
Expanded cohort
Residual Disease
Optimisation cohort
Expanded cohort
Patient Histology
Optimisation cohort
Expanded cohort
Patient cohort comparisons. A. Age density and boxplots by cohort. B. Age density and boxplots by biobank. C. BMI by cohort. D. CA125 by cohort. E. Disease Stage, Surgery Type, Residual Disease, and Patient Histology cohort comparisons.

### Slide 4
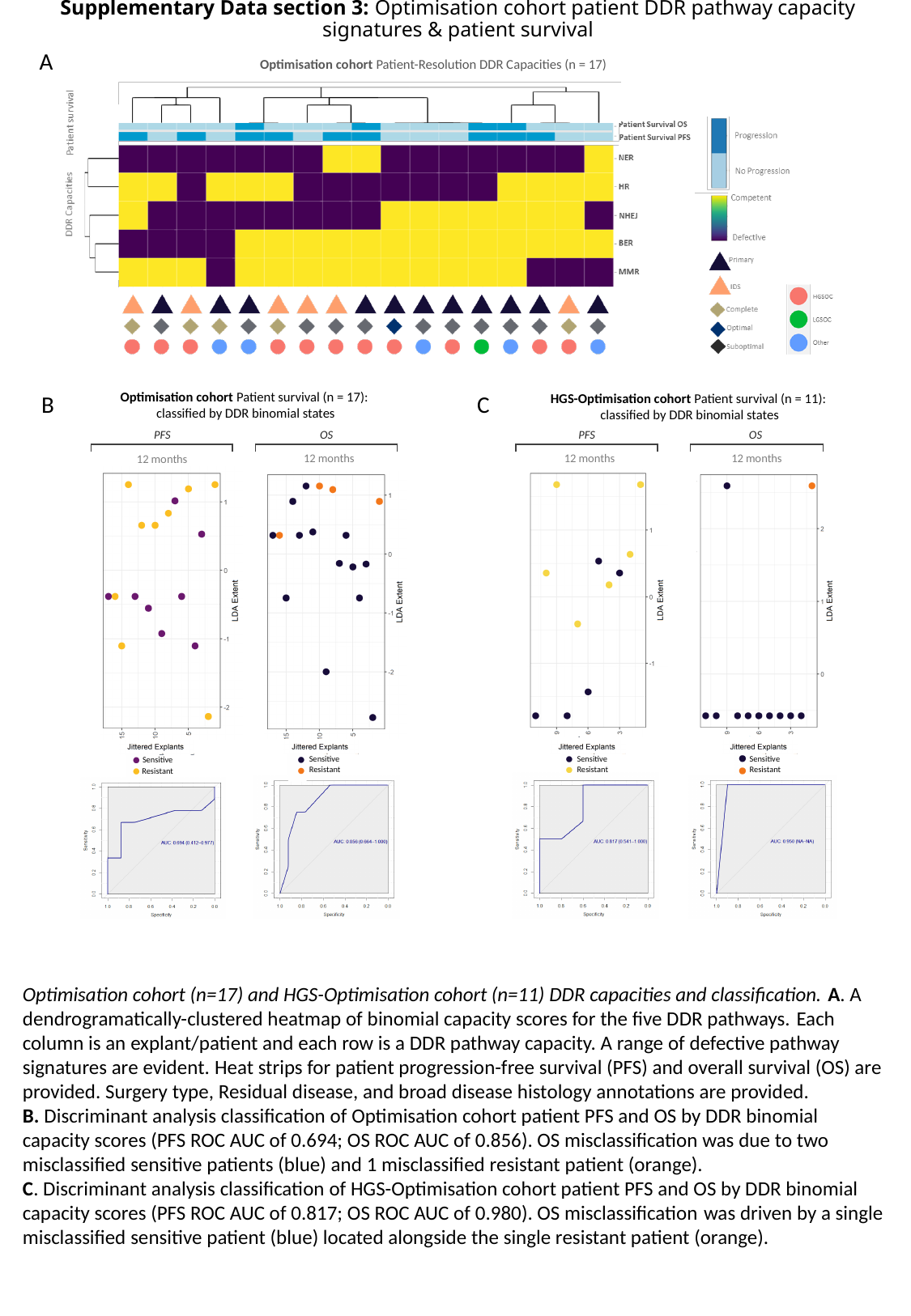

Supplementary Data section 3: Optimisation cohort patient DDR pathway capacity signatures & patient survival
A
Optimisation cohort Patient-Resolution DDR Capacities (n = 17)
Optimisation cohort Patient survival (n = 17):
classified by DDR binomial states
B
C
HGS-Optimisation cohort Patient survival (n = 11):
classified by DDR binomial states
PFS
12 months
PFS
12 months
OS
12 months
OS
12 months
Sensitive
Resistant
Sensitive
Resistant
Sensitive
Resistant
Sensitive
Resistant
Optimisation cohort (n=17) and HGS-Optimisation cohort (n=11) DDR capacities and classification. A. A dendrogramatically-clustered heatmap of binomial capacity scores for the five DDR pathways. Each column is an explant/patient and each row is a DDR pathway capacity. A range of defective pathway signatures are evident. Heat strips for patient progression-free survival (PFS) and overall survival (OS) are provided. Surgery type, Residual disease, and broad disease histology annotations are provided.
B. Discriminant analysis classification of Optimisation cohort patient PFS and OS by DDR binomial capacity scores (PFS ROC AUC of 0.694; OS ROC AUC of 0.856). OS misclassification was due to two misclassified sensitive patients (blue) and 1 misclassified resistant patient (orange).
C. Discriminant analysis classification of HGS-Optimisation cohort patient PFS and OS by DDR binomial capacity scores (PFS ROC AUC of 0.817; OS ROC AUC of 0.980). OS misclassification was driven by a single misclassified sensitive patient (blue) located alongside the single resistant patient (orange).

### Slide 5
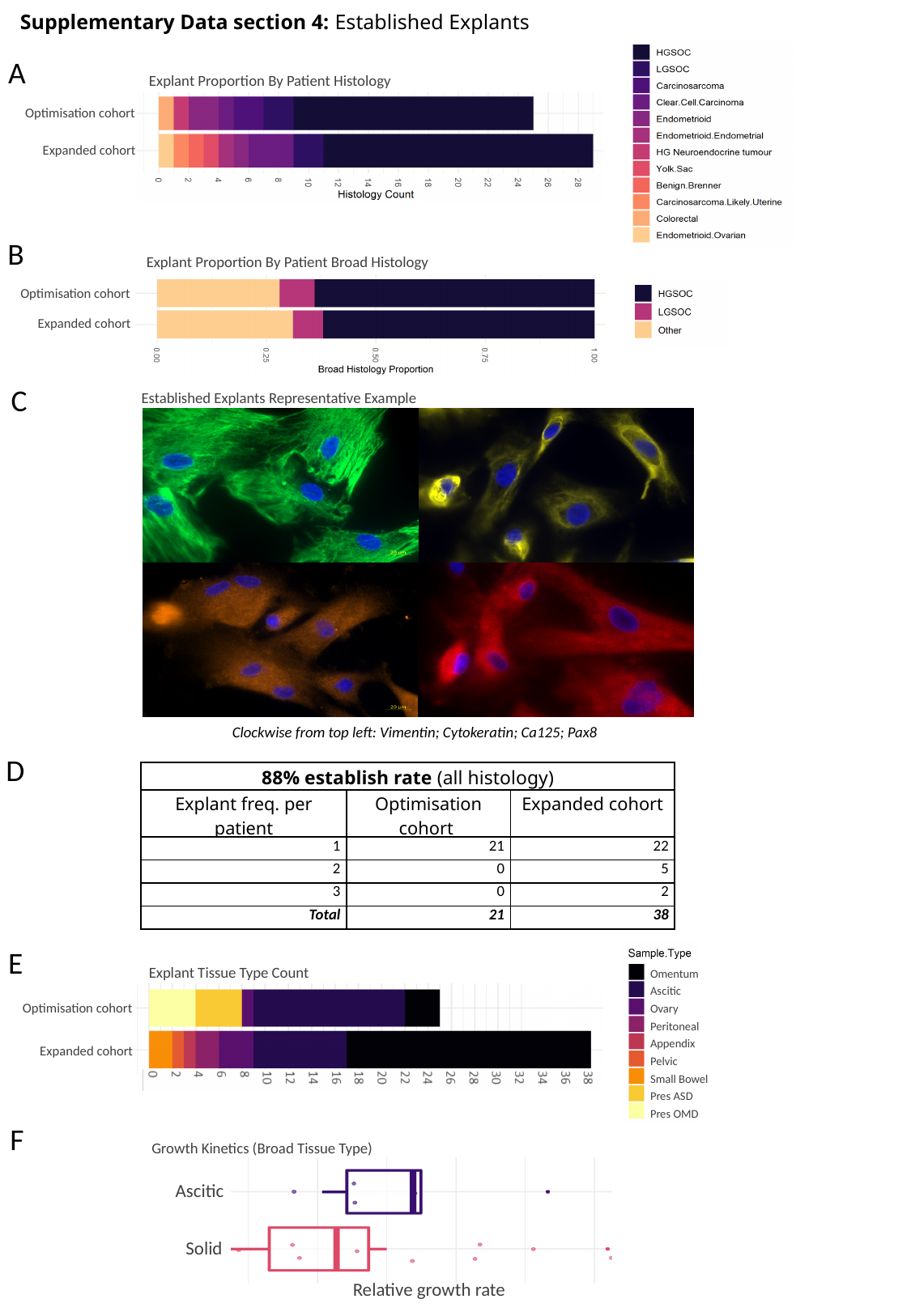

Supplementary Data section 4: Established Explants
A
Explant Proportion By Patient Histology
Optimisation cohort
Expanded cohort
B
Explant Proportion By Patient Broad Histology
Optimisation cohort
Expanded cohort
C
Established Explants Representative Example
Clockwise from top left: Vimentin; Cytokeratin; Ca125; Pax8
D
| 88% establish rate (all histology) | Available samples (all histology) | |
| --- | --- | --- |
| Explant freq. per patient | Optimisation cohort | Expanded cohort |
| 1 | 21 | 22 |
| 2 | 0 | 5 |
| 3 | 0 | 2 |
| Total | 21 | 38 |
 includes NA on some DDR assays
E
Omentum
Ascitic
Ovary
Peritoneal
Appendix
Pelvic
Small Bowel
Pres ASD
Pres OMD
Explant Tissue Type Count
Optimisation cohort
Expanded cohort
F
Growth Kinetics (Broad Tissue Type)
Ascitic
Solid
Relative growth rate

### Slide 6
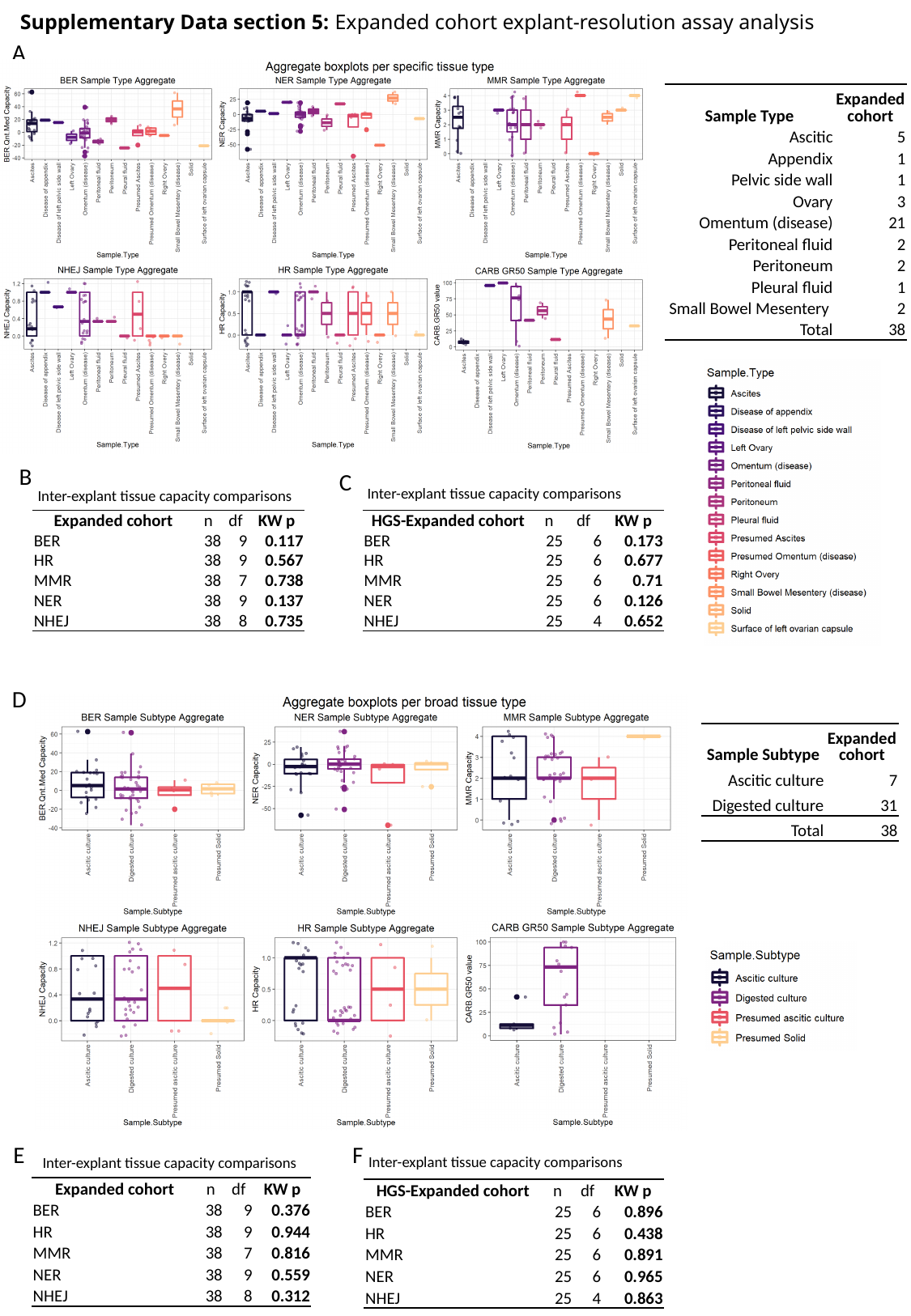

Supplementary Data section 5: Expanded cohort explant-resolution assay analysis
A
| Sample Type | Expanded cohort |
| --- | --- |
| Ascitic | 5 |
| Appendix | 1 |
| Pelvic side wall | 1 |
| Ovary | 3 |
| Omentum (disease) | 21 |
| Peritoneal fluid | 2 |
| Peritoneum | 2 |
| Pleural fluid | 1 |
| Small Bowel Mesentery | 2 |
| Total | 38 |
B
C
Inter-explant tissue capacity comparisons
Inter-explant tissue capacity comparisons
| Expanded cohort | n | df | KW p |
| --- | --- | --- | --- |
| BER | 38 | 9 | 0.117 |
| HR | 38 | 9 | 0.567 |
| MMR | 38 | 7 | 0.738 |
| NER | 38 | 9 | 0.137 |
| NHEJ | 38 | 8 | 0.735 |
| HGS-Expanded cohort | n | df | KW p |
| --- | --- | --- | --- |
| BER | 25 | 6 | 0.173 |
| HR | 25 | 6 | 0.677 |
| MMR | 25 | 6 | 0.71 |
| NER | 25 | 6 | 0.126 |
| NHEJ | 25 | 4 | 0.652 |
D
| Sample Subtype | Expanded cohort |
| --- | --- |
| Ascitic culture | 7 |
| Digested culture | 31 |
| Total | 38 |
E
F
Inter-explant tissue capacity comparisons
Inter-explant tissue capacity comparisons
| Expanded cohort | n | df | KW p |
| --- | --- | --- | --- |
| BER | 38 | 9 | 0.376 |
| HR | 38 | 9 | 0.944 |
| MMR | 38 | 7 | 0.816 |
| NER | 38 | 9 | 0.559 |
| NHEJ | 38 | 8 | 0.312 |
| HGS-Expanded cohort | n | df | KW p |
| --- | --- | --- | --- |
| BER | 25 | 6 | 0.896 |
| HR | 25 | 6 | 0.438 |
| MMR | 25 | 6 | 0.891 |
| NER | 25 | 6 | 0.965 |
| NHEJ | 25 | 4 | 0.863 |

### Slide 7
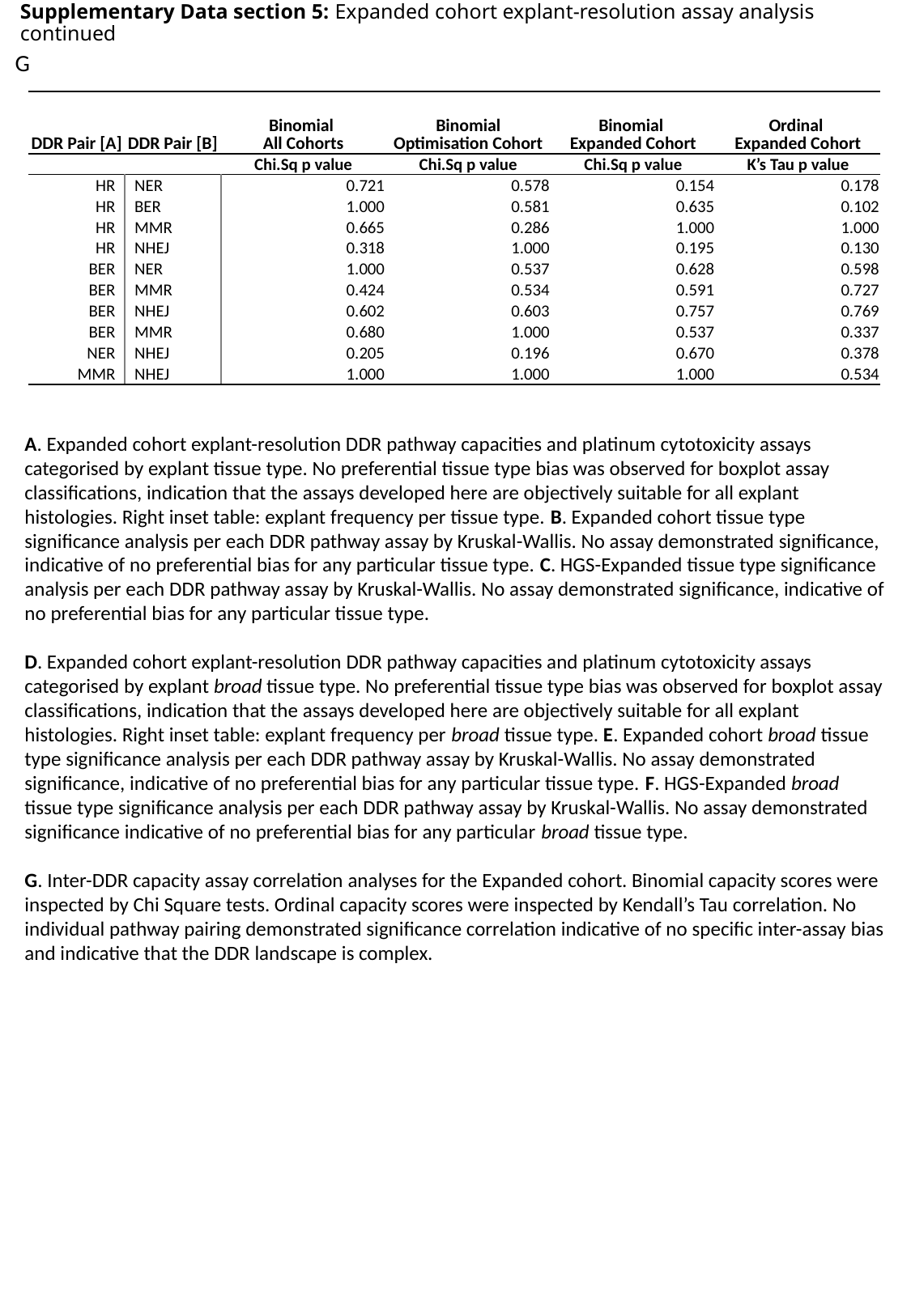

Supplementary Data section 5: Expanded cohort explant-resolution assay analysis continued
G
| DDR Pair [A] | DDR Pair [B] | Binomial All Cohorts | Binomial Optimisation Cohort | Binomial Expanded Cohort | Ordinal Expanded Cohort |
| --- | --- | --- | --- | --- | --- |
| | | Chi.Sq p value | Chi.Sq p value | Chi.Sq p value | K’s Tau p value |
| HR | NER | 0.721 | 0.578 | 0.154 | 0.178 |
| HR | BER | 1.000 | 0.581 | 0.635 | 0.102 |
| HR | MMR | 0.665 | 0.286 | 1.000 | 1.000 |
| HR | NHEJ | 0.318 | 1.000 | 0.195 | 0.130 |
| BER | NER | 1.000 | 0.537 | 0.628 | 0.598 |
| BER | MMR | 0.424 | 0.534 | 0.591 | 0.727 |
| BER | NHEJ | 0.602 | 0.603 | 0.757 | 0.769 |
| BER | MMR | 0.680 | 1.000 | 0.537 | 0.337 |
| NER | NHEJ | 0.205 | 0.196 | 0.670 | 0.378 |
| MMR | NHEJ | 1.000 | 1.000 | 1.000 | 0.534 |
A. Expanded cohort explant-resolution DDR pathway capacities and platinum cytotoxicity assays categorised by explant tissue type. No preferential tissue type bias was observed for boxplot assay classifications, indication that the assays developed here are objectively suitable for all explant histologies. Right inset table: explant frequency per tissue type. B. Expanded cohort tissue type significance analysis per each DDR pathway assay by Kruskal-Wallis. No assay demonstrated significance, indicative of no preferential bias for any particular tissue type. C. HGS-Expanded tissue type significance analysis per each DDR pathway assay by Kruskal-Wallis. No assay demonstrated significance, indicative of no preferential bias for any particular tissue type.
D. Expanded cohort explant-resolution DDR pathway capacities and platinum cytotoxicity assays categorised by explant broad tissue type. No preferential tissue type bias was observed for boxplot assay classifications, indication that the assays developed here are objectively suitable for all explant histologies. Right inset table: explant frequency per broad tissue type. E. Expanded cohort broad tissue type significance analysis per each DDR pathway assay by Kruskal-Wallis. No assay demonstrated significance, indicative of no preferential bias for any particular tissue type. F. HGS-Expanded broad tissue type significance analysis per each DDR pathway assay by Kruskal-Wallis. No assay demonstrated significance indicative of no preferential bias for any particular broad tissue type.
G. Inter-DDR capacity assay correlation analyses for the Expanded cohort. Binomial capacity scores were inspected by Chi Square tests. Ordinal capacity scores were inspected by Kendall’s Tau correlation. No individual pathway pairing demonstrated significance correlation indicative of no specific inter-assay bias and indicative that the DDR landscape is complex.

### Slide 8
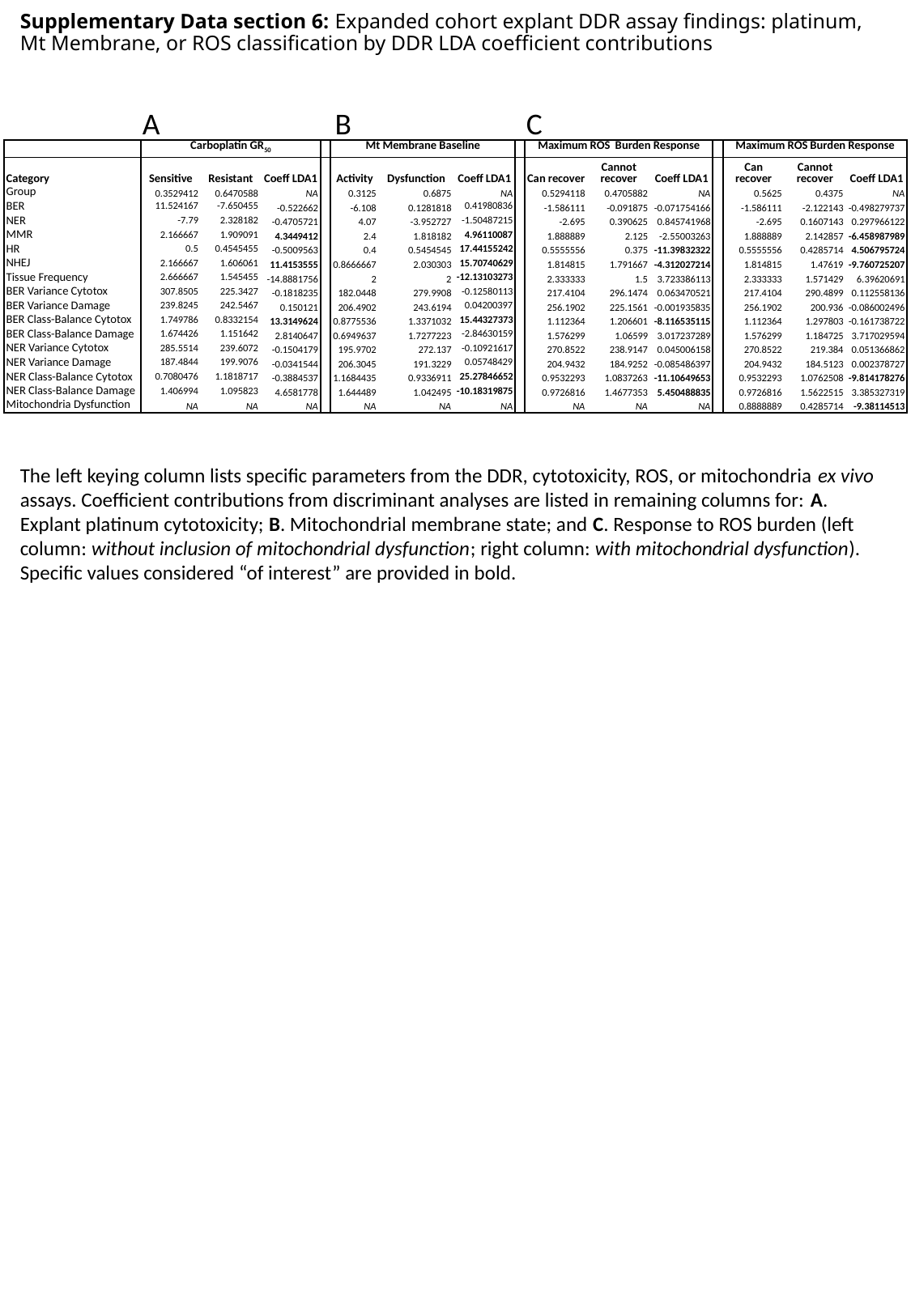

Supplementary Data section 6: Expanded cohort explant DDR assay findings: platinum, Mt Membrane, or ROS classification by DDR LDA coefficient contributions
A
B
C
| | Carboplatin GR_50_ | | | | Mt Membrane Baseline | | | | Maximum ROS Burden Response | | | | Maximum ROS Burden Response | | |
| --- | --- | --- | --- | --- | --- | --- | --- | --- | --- | --- | --- | --- | --- | --- | --- |
| Category | Sensitive | Resistant | Coeff LDA1 | | Activity | Dysfunction | Coeff LDA1 | | Can recover | Cannot recover | Coeff LDA1 | | Can recover | Cannot recover | Coeff LDA1 |
| Group | 0.3529412 | 0.6470588 | NA | | 0.3125 | 0.6875 | NA | | 0.5294118 | 0.4705882 | NA | | 0.5625 | 0.4375 | NA |
| BER | 11.524167 | -7.650455 | -0.522662 | | -6.108 | 0.1281818 | 0.41980836 | | -1.586111 | -0.091875 | -0.071754166 | | -1.586111 | -2.122143 | -0.498279737 |
| NER | -7.79 | 2.328182 | -0.4705721 | | 4.07 | -3.952727 | -1.50487215 | | -2.695 | 0.390625 | 0.845741968 | | -2.695 | 0.1607143 | 0.297966122 |
| MMR | 2.166667 | 1.909091 | 4.3449412 | | 2.4 | 1.818182 | 4.96110087 | | 1.888889 | 2.125 | -2.55003263 | | 1.888889 | 2.142857 | -6.458987989 |
| HR | 0.5 | 0.4545455 | -0.5009563 | | 0.4 | 0.5454545 | 17.44155242 | | 0.5555556 | 0.375 | -11.39832322 | | 0.5555556 | 0.4285714 | 4.506795724 |
| NHEJ | 2.166667 | 1.606061 | 11.4153555 | | 0.8666667 | 2.030303 | 15.70740629 | | 1.814815 | 1.791667 | -4.312027214 | | 1.814815 | 1.47619 | -9.760725207 |
| Tissue Frequency | 2.666667 | 1.545455 | -14.8881756 | | 2 | 2 | -12.13103273 | | 2.333333 | 1.5 | 3.723386113 | | 2.333333 | 1.571429 | 6.39620691 |
| BER Variance Cytotox | 307.8505 | 225.3427 | -0.1818235 | | 182.0448 | 279.9908 | -0.12580113 | | 217.4104 | 296.1474 | 0.063470521 | | 217.4104 | 290.4899 | 0.112558136 |
| BER Variance Damage | 239.8245 | 242.5467 | 0.150121 | | 206.4902 | 243.6194 | 0.04200397 | | 256.1902 | 225.1561 | -0.001935835 | | 256.1902 | 200.936 | -0.086002496 |
| BER Class-Balance Cytotox | 1.749786 | 0.8332154 | 13.3149624 | | 0.8775536 | 1.3371032 | 15.44327373 | | 1.112364 | 1.206601 | -8.116535115 | | 1.112364 | 1.297803 | -0.161738722 |
| BER Class-Balance Damage | 1.674426 | 1.151642 | 2.8140647 | | 0.6949637 | 1.7277223 | -2.84630159 | | 1.576299 | 1.06599 | 3.017237289 | | 1.576299 | 1.184725 | 3.717029594 |
| NER Variance Cytotox | 285.5514 | 239.6072 | -0.1504179 | | 195.9702 | 272.137 | -0.10921617 | | 270.8522 | 238.9147 | 0.045006158 | | 270.8522 | 219.384 | 0.051366862 |
| NER Variance Damage | 187.4844 | 199.9076 | -0.0341544 | | 206.3045 | 191.3229 | 0.05748429 | | 204.9432 | 184.9252 | -0.085486397 | | 204.9432 | 184.5123 | 0.002378727 |
| NER Class-Balance Cytotox | 0.7080476 | 1.1818717 | -0.3884537 | | 1.1684435 | 0.9336911 | 25.27846652 | | 0.9532293 | 1.0837263 | -11.10649653 | | 0.9532293 | 1.0762508 | -9.814178276 |
| NER Class-Balance Damage | 1.406994 | 1.095823 | 4.6581778 | | 1.644489 | 1.042495 | -10.18319875 | | 0.9726816 | 1.4677353 | 5.450488835 | | 0.9726816 | 1.5622515 | 3.385327319 |
| Mitochondria Dysfunction | NA | NA | NA | | NA | NA | NA | | NA | NA | NA | | 0.8888889 | 0.4285714 | -9.38114513 |
The left keying column lists specific parameters from the DDR, cytotoxicity, ROS, or mitochondria ex vivo assays. Coefficient contributions from discriminant analyses are listed in remaining columns for: A. Explant platinum cytotoxicity; B. Mitochondrial membrane state; and C. Response to ROS burden (left column: without inclusion of mitochondrial dysfunction; right column: with mitochondrial dysfunction). Specific values considered “of interest” are provided in bold.

### Slide 9
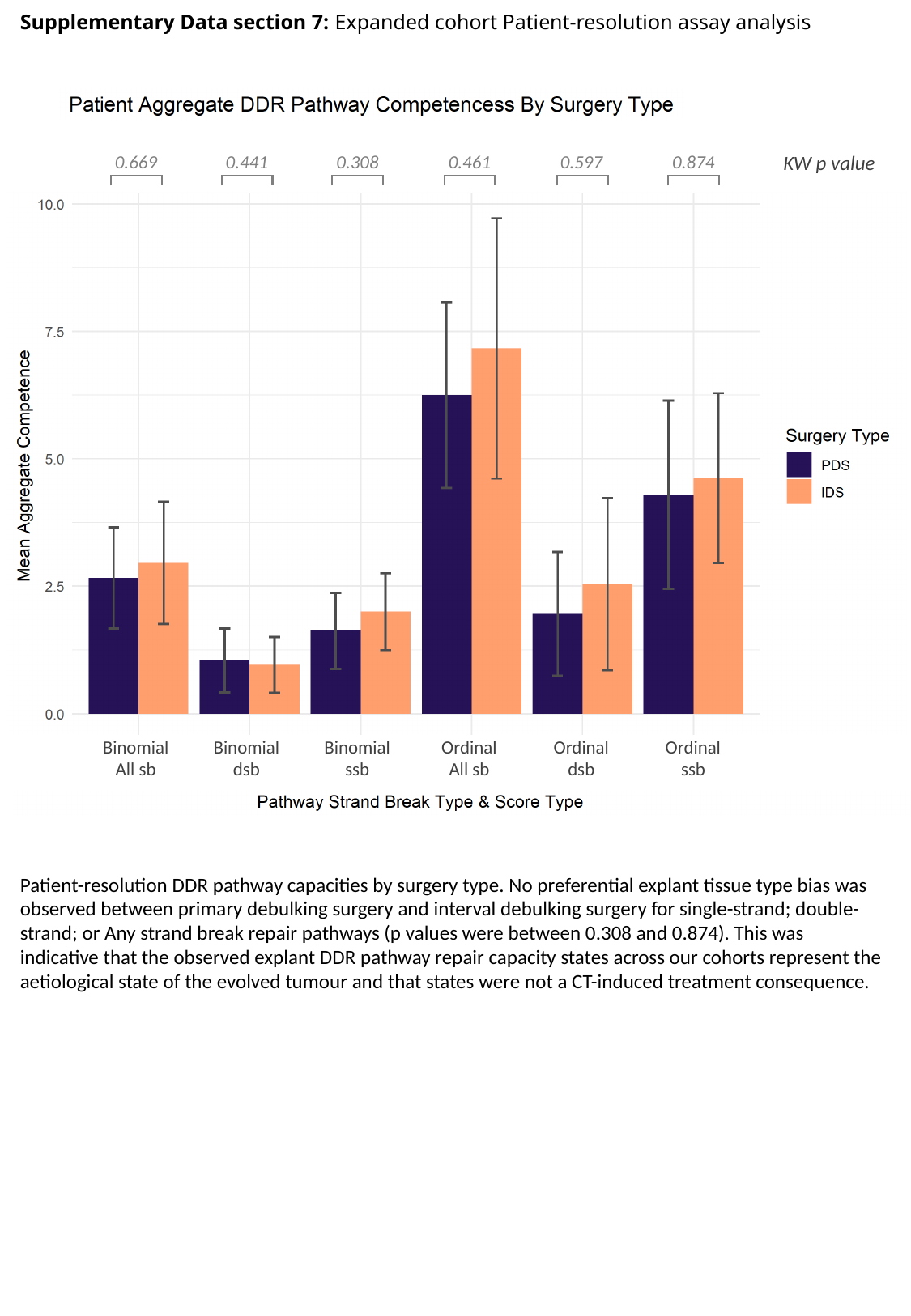

Supplementary Data section 7: Expanded cohort Patient-resolution assay analysis
KW p value
0.669
0.441
0.308
0.461
0.597
0.874
Binomial
All sb
Binomial
dsb
Binomial
ssb
Ordinal
All sb
Ordinal
dsb
Ordinal
ssb
Patient-resolution DDR pathway capacities by surgery type. No preferential explant tissue type bias was observed between primary debulking surgery and interval debulking surgery for single-strand; double-strand; or Any strand break repair pathways (p values were between 0.308 and 0.874). This was indicative that the observed explant DDR pathway repair capacity states across our cohorts represent the aetiological state of the evolved tumour and that states were not a CT-induced treatment consequence.
